# Astroglial exosome HepaCAM signaling and ApoE antagonization coordinates early postnatal cortical pyramidal neuronal axon growth and dendritic spine formation

**DOI:** 10.1101/2023.02.14.528554

**Authors:** Shijie Jin, Xuan Chen, Yang Tian, Rachel Jarvis, Vanessa Promes, Yongjie Yang

**Author notes:** To whom correspondence and material request should be addressed: Yongjie Yang, Tufts University, Department of Neuroscience, 136 Harrison Ave, Boston, MA 02111, USA.

## Abstract

Developing astroglia play important roles in regulating synaptogenesis through secreted and contact signals. Whether they regulate postnatal axon growth is unknown. By selectively isolating exosomes using size-exclusion chromatography (SEC) and employing cell-type specific exosome reporter mice, our current results define a secreted astroglial exosome pathway that can spread long-range *in vivo* and stimulate axon growth of cortical pyramidal neurons. Subsequent biochemical and genetic studies found that surface expression of glial HepaCAM protein essentially and sufficiently mediates the axon-stimulating effect of astroglial exosomes. Interestingly, apolipoprotein E (ApoE), a major astroglia-secreted cholesterol carrier to promote synaptogenesis, strongly inhibits the stimulatory effect of astroglial exosomes on axon growth. Developmental ApoE deficiency also significantly reduces spine density of cortical pyramidal neurons. Together, our study suggests a surface contact mechanism of astroglial exosomes in regulating axon growth and its antagonization by ApoE, which collectively coordinates early postnatal pyramidal neuronal axon growth and dendritic spine formation.

## Introduction

Developmental neuronal axon outgrowth and synaptogenesis are crucial steps in forming sophisticated and functional connectivity in the mammalian central nervous system (CNS). It is well established that synaptogenesis begins after birth and continues for several weeks postnatally while axon outgrowth is mostly completed at birth^1^. However, descending corticospinal tract (CST) axons that predominantly originate from layer V pyramidal neurons of the primary motor cortex continue to grow and reach spinal cord segments from postnatal day 1 to 10 (P1 to P10) in mice^2^. Developing astroglia have been well demonstrated to actively promote synaptogenesis and synapse maturation^3^. Early studies established that glia-derived cholesterol, transported by the astroglia-secreted lipoprotein ApoE, serves as a robust synaptogenic factor for retinal ganglion cells (RGCs)^4^. Several other secreted proteins from astroglia, such as Thrombospondin 1 and 2 (Tsp1/2)^5^, Hevin^6^, glypicans^7^, and Chordin-like 1^8^, have been later identified to promote excitatory synapse formation and stimulate glutamatergic activity. In contrast, far less is understood about whether and how developing astroglia regulate axon growth.

Developmental axon growth is driven by the actin and microtube dynamics within axonal growth cones as a result of receptor activation by extracellular trophic factors, adhesion molecules, and matrix proteins^9^. Although many of these ECM/adhesion proteins, such as neural cell adhesion molecule (NCAM), N-cadherin, and integrins, are highly expressed in developing neurons and neural progenitors^10^, transcriptome profiling has found expression of a number of ECM and CAM genes in developing astroglia^11^. Early studies showed that γ-protocadherins (γ-Pcdhs) are also expressed by astroglia which promotes synaptogenesis *in vitro* and *in vivo*^12^. Genetic studies found that astroglial expression of neuroligins is important for developmental astroglial morphogenesis^13^. Neuronal cell adhesion molecule (NrCAM) was also found at astroglial process to regulate astroglia-inhibitory synapse interaction^14^. In particular, hepatocyte cell adhesion molecule (HepaCAM, also known as GlialCAM), a CAM protein containing immunoglobulin (Ig)-like extracellular domains^15^, is highly enriched in (astro)glia in the CNS^16^. HepaCAM has been identified as a binding partner for a voltage-gated chloride channel Clc-2 and its mutations have been implicated in causing a rare form of leukodystrophy^17, 18^. HepaCAM was also recently shown to regulate astroglial domain territory and gap junction coupling^19^. Whether astroglial CAM proteins including HepaCAM play a role in developmental axon growth remains unknown.

Exosomes (50-150 nm in diameter), a major type of secreted extracellular vesicles (EVs), are derived from intraluminal vesicles (ILVs) in the early endosomal compartment and are released from multivesicular bodies (MVBs) during endosome maturation^20^. EVs and exosomes secreted from various CNS cell types have been shown to regulate activity-dependent translation^21^ and glutamate transporter function^22^, to promote axon myelination and transport^23^, and to maintain brain vascular integrity^24^. Whether astroglial exosome signals play a role in regulating neuronal functions has just begun to be understood. Astroglia derived extracellular vesicles (ADEVs) are able to modulate dendritic complexity of cultured hippocampal neurons^25^. An extracellular matrix protein, fibulin-2, was also recently identified as astrocyte EV cargo that promotes synapse formation in a TGFβ-dependent manner^26^. However, the ultracentrifugation (UC) approach used in these studies to isolate astroglial exosomes often leads to mixed exosomes and secreted proteins^27^, potentially undermining the effects mediated by astroglia secreted exosomes.

In the current study, we investigated the developmental function of astroglial exosomes, especially surface HepaCAM signaling, in regulating axon growth of cortical pyramidal neurons and how this pathway is antagonized by ApoE, which collectively coordinates early postnatal pyramidal neuronal axon growth and dendritic spine formation.

## Results

### Size exclusion chromatography (SEC)-isolated astroglial exosomes (A-Exo.) stimulate axon growth of cortical neurons

Exosomes have been conventionally isolated from cell culture medium or body fluids using serial (ultra)centrifugation steps^28^. However, recent studies have shown that UC-isolated exosomes are often contaminated with secreted proteins from cells^27, 29^. We initially isolated A-Exo. from astrocyte conditioned medium (ACM, conditioned from > 90% confluent astrocytes for 3d) using the UC method and detected well-validated exosome markers, including tetraspanin family proteins CD63/CD81 and the ESCRT protein Tsg101^20^, together with several astroglia-secreted proteins, such as Tsp1/2, Hevin, Sema3A, and Sparc (Supplementary Fig. 1a) that were previously identified as astroglia-secreted synaptogenesis modulators^3^. To better separate A-Exo. from secreted proteins, we optimized exosome isolation procedures using filtration (0.22μm) and SEC (Fig. 1a). Immunoblotting of astroglia-secreted proteins and exosome markers in representative eluted fractions from ACM showed that Tsp1/2, Sema3A, Hevin, and Sparc are only detected in exosome-free but not in CD81^+^ A-Exo. fractions of ACM (Fig. 1b). ImmunoEM analysis of CD63 in eluted fractions further confirmed that CD63^+^ exosomal vesicles are detected only in exosome (#7-8) and mixed (#9) fractions (white arrows, Fig. 1c ii-iii) but not in other ACM fractions (Fig. 1c, i, iv; Supplementary Fig. 1b). Notably, translucent CD63^-^ small vesicles (30-40 nm size range), possibly exomeres^30^, were observed in certain eluted fractions, especially in exosome-free ACM fractions (yellow arrows, Fig. 1c ii-iii; Supplementary Fig. 1b). Exosome fractions were also analyzed by the qNano particle analyzer^31^ from which a single Gaussian peak at a mean of 70-80 nm (Supplementary Fig. 1c) was revealed, confirming the population of nanovesicles with the typical size of exosomes.

**Fig. 1.**
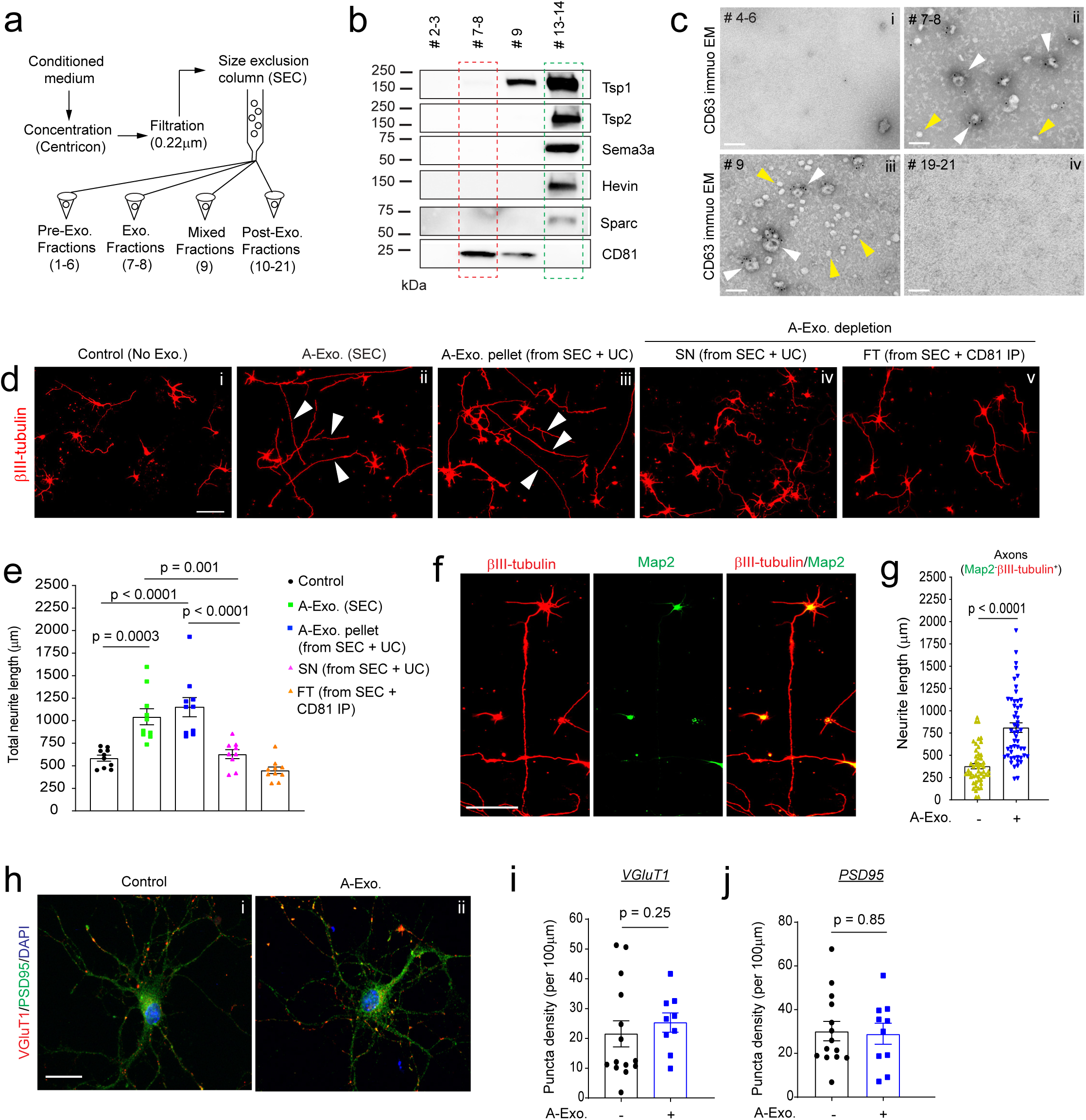
Size exclusion chromatography (SEC)-isolated astroglial exosomes (A-Exo.) selectively stimulate neuronal axon growth. **a,** Schematic diagram of SEC-based isolation of exosomes from ACM. A 10k molecular weight cutoff Centricon® Plus-70 centrifugal filter device was used; **b,** Representative immunoblots of astroglia secreted proteins and exosome marker CD81 from eluted fractions (pooled as indicated, 500 μl/fraction) of ACM (100 mL/sample) from SEC. Unconcentrated elution (10 μl/sample) was run on immunoblot; **c,** Representative immunoEM images of CD63 labeling in different SEC eluted fractions. Subpanels i-iv: fractions #4-6, #7-8, #9, and #19-21, respectively; white arrows: CD63^+^ A-Exo.; yellow arrows: CD63^-^ small vesicles; scale bar: 100 nm. Representative images (**d**) and quantification (**e**) of βIII-tubulin^+^ neurite length of cortical neurons in control (i) or treated with SEC-isolated A-Exo. (ii), A-Exo. pellet (iii, from SEC + UC), A-Exo. depleted SN (iv, from SEC +UC), or FT (v, from SEC + CD81 IP); white arrows: elongated neurites; scale bar 100 µm. n = 10 neurons (2 biological replicates)/group; **f**, Representative image of βIII-tubulin and Map2 stained cortical neurons following A-Exo. treatment. Scale bar: 100 μm; **g**, Quantification of Map2^-^βIII-tubulin^+^ axon length following A-Exo. treatment. n = 51-55 neurons (> 3 biological replicates)/group; Representative image of VGluT1 and PSD95 staining on cortical neuronal cultures (**h**) and quantification of VGluT1 density (**i**) and PSD95 density (**j**) n = 10-14 neurons (2 biological replicates)/group; p values in **e** determined by one-way ANOVA followed by post-hoc Tukey’s test; p values in **g**, **i**, and **j** determined by two-tailed t test.

Whether and how A-Exo. influence neuronal properties is little known. Although tetraspanin protein (CD63 or CD81) immunoprecipitation (IP) can selectively isolate A-Exo., removing exosomes from IP beads has been difficult and the wash solution often kills neurons. As an alternative, we directly treated cultured cortical neurons with SEC-eluted pre-exosome (#4-6), exosome (#7-8), and post-exosome (#10-12 and #19-21 respectively) fractions from ACM for 24hr. Interestingly, βIII-tubulin^+^ neurites from neuronal cultures treated (at DIV 4) with exosome fractions, but not other fractions, are substantially longer than in the untreated control (Supplementary Fig. 2a-b). A-Exo.-stimulated neurite growth is also treatment time-dependent, with < 10% or > 50% of neurites longer than 600 μm after either 1 or 3d treatment, respectively (Supplementary Fig. 2c). In contrast, HEK cell-secreted exosomes have no stimulating effect on neurite growth (Supplementary Fig. 2d), indicating a specific effect of A-Exo. on neurite growth. As we observed CD63^-^ translucent vesicles in exosome fractions (Fig. 1c) from the SEC procedure, to confirm that A-Exo. indeed stimulates neurite growth, exosomes were depleted from SEC-eluted exosome fractions by CD81 IP or by an additional UC step (100,000 x g for 24h). Both CD81 IP and the additional UC step effectively depleted exosomes, indicated by the detection of CD81 expression only in CD81 IP and UC pellets but not in flow-through (FT) from CD81 IP or in supernatant (SN) from the UC step (Supplementary Fig. 2e). Consistently, exosome-depleted FT from CD81 IP or SN from the additional UC step has no effect on stimulating neurite growth (Fig. 1d iv-v, Fig. 1e), while the pelleted A-Exo. from the additional UC step retain the stimulatory effect on neurite growth (Fig. 1d iii, Fig. 1e).

Subsequent immunostaining of neurite markers indicates that axons (Map2^-^βIII-tubulin^+^) but not dendrites (Map2^+^βIII-tubulin^+^) are specifically elongated by A-Exo. treatment (Fig. 1f-g, Supplementary Fig. 2f). Active axonal elongation of cortical neurons induced by A-Exo. was also observed in time-lapse live cell imaging (8h time frame, Supplementary Movie). Immunostaining of additional axon markers such as Tau was also performed to confirm axon-specific stimulation by A-Exo. (Supplementary Fig. 2g). βIII-tubulin staining was then primarily shown for neurite labeling in subsequent results. Interestingly, A-Exo. treatment induces no significant changes in neuronal morphology and synapse numbers (Fig. 1h), indicated by similar neurite VGluT1 and PSD95 density (quantified from secondary branches, Fig. 1 i-j). Consistently, Sholl analysis of cortical neurons also confirmed that the overall morphological complexity of cortical neurons is not altered by A-Exo. treatment, other than continuous intersections at distal but not proximal (< 150 μm) distances from the soma (Supplementary Fig. 2h), as a result of elongated axons.

### Surface expression of HepaCAM (GlialCAM) mediates stimulatory effect of A-Exo. on axon growth

To begin dissecting how A-Exo. stimulate axon growth, we performed different biochemical treatments, i.e., proteinase K, RNase, and/or sonication on A-Exo., to examine whether RNA or proteins especially surface proteins, mediate the stimulatory effect of A-Exo. on axon growth. To test whether RNA (including microRNA) in exosomes is involved in exosome-mediated stimulation of axon growth, sonicated and RNase treated A-Exo. (1 μg/sample) were added onto cortical neuronal cultures. Interestingly, exosomes with essentially all RNA degraded, as confirmed by bioanalyzer analysis (Supplementary Fig. 3a), are still able to strongly stimulate neurite growth, similarly to untreated A-Exo. (White arrows, Fig. 2a vi, Fig. 2b), supporting the non-involvement of RNA in mediating the stimulatory effect of A-Exo. on axon growth. In contrast, proteinase K treatment of A-Exo. in which surface exosomal proteins, such as CD81, are degraded (Supplementary Fig. 3b) completely abolished the stimulatory effect of A-Exo. on axon growth (Fig. 2a ii-iii, Fig. 2b). In addition, sonicated A-Exo. surface fractions without lysate remain equally as stimulatory as untreated A-Exo. (White arrows, Fig. 2a iv-v, Fig. 2b). These results point to a potential surface protein mechanism in mediating the stimulatory effect of A-Exo. on axon growth. We further tested the involvement of A-Exo. surface contact with neurons by plating cortical neurons onto coverslips that were coated with poly-D-lysine (PDL), PDL/laminin (LN), PDL + A-Exo., or PDL/LN + A-Exo. Consistent with the results from biochemical treatments of A-Exo., neuronal axons are significantly longer on PDL or PDL/LN with A-Exo. -coated coverslips compared to PDL or PDL/LN coated alone (Fig. 2c-d, Supplementary Fig. 3c). Additionally, inhibition of clathrin-dependent endocytosis by dynasore, a cell-permeable inhibitor of dynamin^32^, has no effect on A-Exo. -stimulated neuronal axon growth (Fig. 2e), excluding the possibility of clathrin-mediated endocytosis of A-Exo. in promoting neuronal axon growth. Together, these results support the notion that surface protein-mediated contact mechanisms mediate the axon-stimulating effect of A-Exo.

**Fig. 2.**
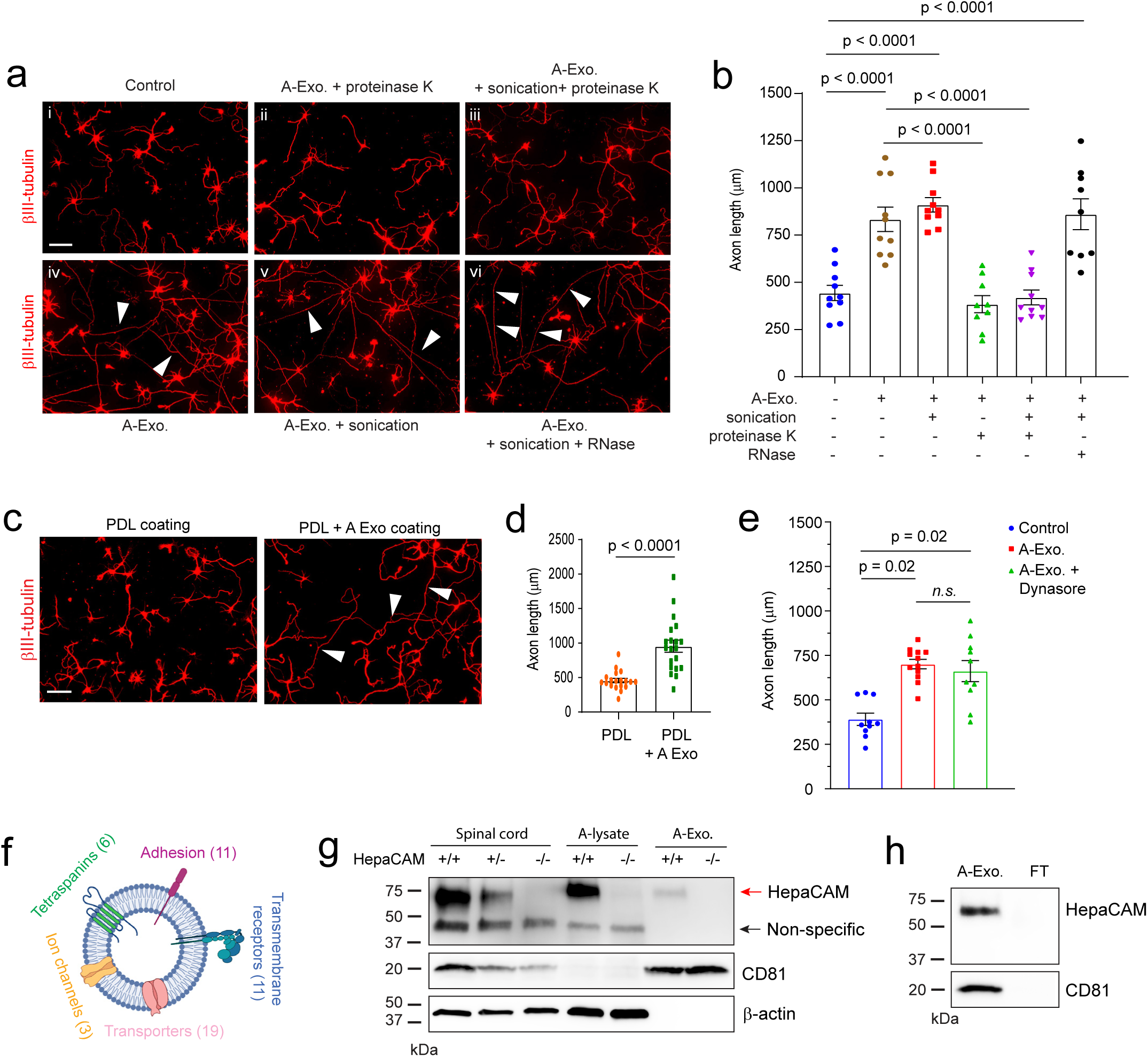
Involvement of A-Exo. surface signals in promoting axon growth and identification of the surface expression of HepaCAM (GlialCAM) on A-Exo. Representative images (**a**) and quantification (**b**) of axon length of cortical neurons in control (i) or treated with proteinase K (10 μg/mL, 5 minutes) digested A-Exo. (ii), sonicated (30s) and proteinase K digested A-Exo. (iii), A-Exo. (iv), sonicated A-Exo. (v), or sonicated (30s) and RNase (10 μg/mL, 5 minutes) digested A-Exo. (vi). 1 μg A-Exo./sample was used in each treatment in **a-b.** White arrows: elongated axons; n = 9-10 neurons (2 biological replicates)/group; Scale bar: 100 μm; Representative images (**c**) and quantification (**d**) of axon length of cortical neurons plated on either poly-D-lysine (PDL) coated or PDL/A-Exo. coated coverslips. n = 20 neurons (2 biological replicates)/group; Scale bar: 100 μm; **e**, Quantification of axon length of cortical neurons following A-Exo. treatment or co-treatment with A-Exo. and dynasore (dynamin inhibitor, 50 μM). n = 10 neurons (2 biological replicates)/group; **f**, Proteomic identification of different categories of transmembrane proteins on A-Exo. surface. Specific transmembrane proteins are included in the Supplementary Table 1. n = 3 biological replicates; **g**, Detection of specific HepaCAM immunoreactivity from spinal cord lysate (10 μg/lane), astrocyte lysate (10 μg/lane), and A-Exo. (1 μg/lane) prepared from WT (+/+), HepaCAM heterozygous (+/-), and HepaCAM KO (-/-) mice; Red arrow: specific HepaCAM immunoreactivity; Black arrow: non-specific immunoreactivity; **h**, Detection of specific HepaCAM immunoreactivity in A-Exo. but not in exosome-free ACM fractions; 1 μg A-Exo. was used in each experiment. p value in **d** determined from two-tailed t test; p values in **b** and **e** determined by one-way ANOVA followed by post-hoc Tukey’s test.

Surface proteins (and internal protein cargos) of A-Exo. remain essentially unknown. As molecular cargoes in exosomes are highly heterogeneous and cell-type dependent ^20^, we performed proteomic analysis on A-Exo. by in-gel trypsin digestion and LC/MS/MS analysis. A total of 347 proteins were identified based on 3 peptides detected per protein and iBAQ > 1 x 10^5^. We used Ingenuity Pathway Analysis (IPA) to specifically analyze transmembrane proteins detected on A-Exo. and found tetraspanins (exosome markers), cell-adhesion molecules (CAMs), transmembrane receptors, transporters, and channels (Fig. 2f, Supplementary Table 1). In particular, HepaCAM (also named GlialCAM), a transmembrane CAM protein highly enriched in CNS astroglia^16^, was found on the surface of A-Exo. Specific HepaCAM immunoreactivity (∼70 KDa size) was also determined and verified in spinal cord, astrocyte lysate, and A-Exo. samples from WT (+/+), HepaCAM heterozygous (+/-), and KO (-/-) mice (generated from HepaCAM floxed mice)^19^ (Fig. 2g). Additionally, HepaCAM was detected only in A-Exo. but not in non-exosome FT in ACM (Fig. 2h), consistent with its characterization as a transmembrane protein. Although the naïve form of HepaCAM protein is predicted to be ∼50 KDa, its glycosylated and membrane associated form has been detected at ∼70 KDa as shown here and previously^16^. Thus, the detection of the glycosylated but not the naïve form of HepaCAM in exosome samples (Fig. 2g-h) also supports the functional role of HepaCAM on exosomal surface. In addition, although certain transmembrane proteins undergo proteolytic cleavage to release their extracellular domain (ECD)^33^, we found no specific HepaCAM immunoreactivity band (∼40 KDa) that would correspond with the size of cleaved ECD in our HepaCAM immunoblots (Supplementary Fig. 3d, Fig. 2g-h), ruling out the possibility that HepaCAM undergoes proteolytic cleavage to release its ECD *in vitro* and *in vivo*.

Although HepaCAM belongs to the CAM family with Ig-like extracellular domains ^15^, its involvement in axon growth remains unexplored. We tested whether HepaCAM is involved in mediating the stimulatory effect of A-Exo. on axon growth by treating cortical neurons with HepaCAM-depleted A-Exo., prepared from HepaCAM KO mouse astrocyte cultures. As shown in Fig. 3a, equal amount (1μg) of HepaCAM-depleted A-Exo. only modestly stimulate neurite growth compared to WT A-Exo. (a 45% reduction, p < 0.0001, Fig. 3b), demonstrating the essential role of HepaCAM in mediating the axon-stimulating effect of A-Exo. This is consistent with the observation that HEK cell exosomes, which do not stimulate axon growth (Supplementary Fig. 2d), lack HepaCAM expression (Supplementary Fig. 3e). Previous studies have shown that HepaCAM depletion dysregulates proper targeting of surface proteins such as Mlc1 and the chloride channel Clc-2 in glial cells^34^. To further demonstrate that the axon-stimulating effect of A-Exo. is mediated directly by HepaCAM but is not due to mistargeted surface proteins resulting from the HepaCAM depletion, we treated cortical neurons with both HepaCAM antibody and A-Exo. The addition of HepaCAM antibody effectively and completely blocked A-Exo’s stimulatory effect on axon growth (Fig. 3c iv, Fig. 3d), while the control IgG antibody had no effect (Fig. 3c iii) on A-Exo’s stimulation of axon growth. IgG itself also has no effect on neuronal axon growth (Fig. 3c ii). To further demonstrate that HepaCAM is sufficient to stimulate axon growth, coverslips were directly coated with PDL and either the HepaCAM extracellular domain (ECD) or BSA. HepaCAM ECD, but not BSA, sufficiently and significantly stimulates axon growth (Fig. 3e iii, Fig. 3f). Together, these genetic and biochemical analyses clearly support the direct role of HepaCAM ECD in mediating the axon-stimulating effect of A-Exo.

**Fig. 3.**
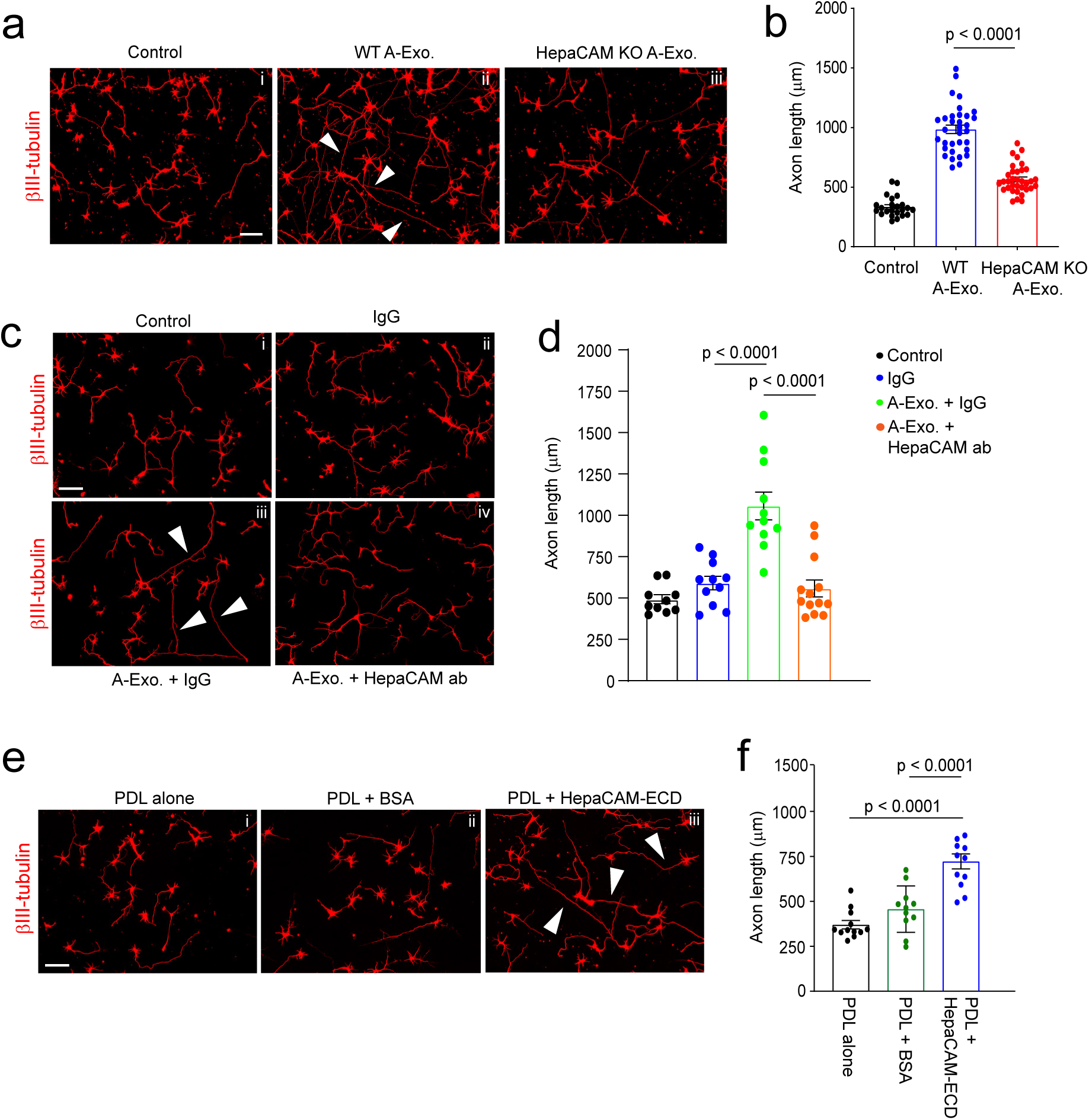
Surface expression of HepaCAM essentially and sufficiently mediates stimulatory effects of A-Exo. on axon growth. Representative images (**a**) and quantification (**b**) of βIII-tubulin^+^ neuronal axon (white arrows) length following equal amount (1μg) of WT and HepaCAM-depleted A-Exo. treatment. Subpanels: i, control; ii, WT A-Exo.; iii, HepaCAM KO A-Exo.; HepaCAM-depleted A-Exo. were prepared from HepaCAM KO astrocyte cultures as described in materials and methods. Scale bar: 100 μm; n = 23-34 neurons (> 3 biological replicates)/group; Representative images (**c**) and quantification (**d**) of βIII-tubulin^+^ neuronal axon (white arrows) length following co-treatment with HepaCAM antibody (ProteinTech) and A-Exo. Subpanels: i, control (1 x PBS); ii, IgG alone; iii, A-Exo. + IgG; iv, A-Exo. + HepaCAM ab; 8 μg ab/coverslip (12 mm diameter) was used in the treatment. Scale bar: 100 μm; n=10-14 neurons (≥ 2 biological replicates)/group; Representative images (**e**) and quantification (**f**) of βIII-tubulin^+^ neuronal axon (white arrows) length following HepaCAM ECD coating. Subpanels: i, PDL alone; ii, PDL + BSA (4 μg); iii, PDL + HepaCAM ECD (4 μg); Scale bar: 100 μm; n=12 neurons (≥ 2 biological replicates)/group; 1μg A-Exo. was used in each experiment; p values in **b**, **d**, and **f** determined using one-way ANOVA followed by a Tukey post-hoc test.

### Developmental dynamics and *in situ* distribution of astroglial exosomes in the CNS

Although a number of *in vitro* studies have reported secretion of exosomes from cultured astroglia, *in situ* distribution and developmental changes of A-Exo. in the CNS remain unexplored, primarily due to the difficulty of selectively labeling cell-type specific exosomes by immunostaining. We previously generated cell-type specific exosome reporter CD63-GFP^f/f^ mice^35^, which allows labeling of cell-type specific exosomes and their intracellular precursors, intraluminal vesicles (ILVs) and multiple vesicular bodies (MVBs). By employing this mouse tool and confocal/immunoEM imaging, we have previously characterized neuronal ILVs and exosomes *in situ* in the CNS^35^. To determine the *in vivo* distribution of A-Exo. in the developing CNS, we generated CD63-GFP^f/+^Ai14-tdT^f/+^ mice and performed stereotaxic injections of AAV5-mCherry-*Gfap*-Cre (0.3μl, 4 x 10^12^ gc/mL) on the motor cortex at either P1 or P21 for tissue collection at P8 or P28, respectively (Fig. 4a). This combined AAV5-mCherry-*Gfap*-Cre and CD63-GFP^f/+^Ai14-tdT^f/+^ mice paradigm allows selective labeling of both astroglial morphology and astroglia secreted exosomes (as well as ILVs/MVBs) simultaneously, which facilitates identification of secreted astroglial exosomes from the same labeled astroglia. CD63-GFP^+^ puncta were found to be clearly co-localized with tdT^+^ astroglial soma and processes at both P8 and P28 (Fig. 4b, Supplementary Fig. 4a). By converting confocal images (Fig. 4b i, iii) into 3D images (Fig. 4b ii, iv) using Imaris image analysis software and quantifying extracellular (secreted exosomes, yellow arrows, Fig. 4b i-iv) and intracellular (ILVs/MVBs, white arrows, Fig. 4b iii) CD63-GFP^+^ puncta based on tdT^+^ astroglial labeling, significantly more CD63-GFP^+^ puncta were observed outside of tdT^+^ cortical astroglia at P8 (55.1%) when astroglial processes are largely undeveloped^13, 36^ than P28 (34.4%) when astroglial processes are fully developed (Fig. 4c), suggesting that astroglial exosomes are particularly and abundantly secreted during first postnatal week when astroglial processes are still primitive.

**Fig. 4.**
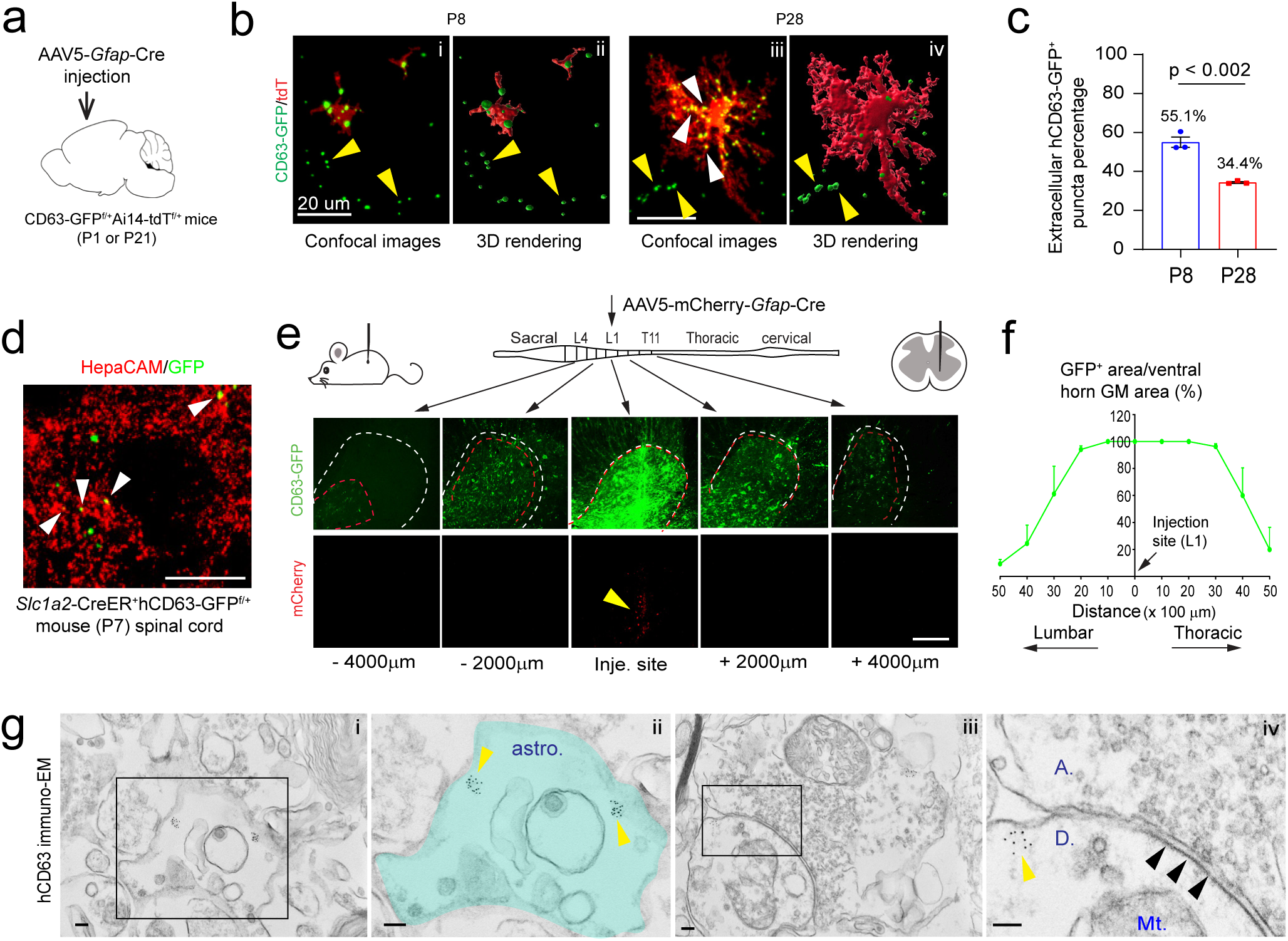
*In situ* illustration and developmental dynamics of A-Exo. in the CNS. **a,** Schematic diagram of stereotaxic injections of AAV5-mCherry-*Gfap*-Cre into the motor cortex of CD63-GFP^f/+^Ai14-tdT^f/f^ mice at P1 or P21. Mice were collected for analysis at P8 or P28, respectively. **b,** Representative confocal and Imaris images of tdT^+^ astroglia and CD63-GFP^+^ puncta at P8 and P28 in AAV5-mCherry-*Gfap*-Cre-injected CD63-GFP^f/+^Ai14-tdT^f/+^ mice. Yellow and white arrows indicate extracellularly or intracellularly localized CD63-GFP^+^ puncta, respectively, based on their co-localization with tdT^+^ astroglia; **c,** Quantification of extracellularly localized CD63-GFP^+^ puncta based on their co-localization with tdT^+^ astroglia; n = 9 images from 3 mice/group; **d**, Representative image of HepaCAM immunostaining signals co-localized with CD63-GFP^+^ puncta signals from spinal cord sections of 4-OHT-injected *Slc1a3*-CreER^+^CD63-GFP^f/+^ mice (P7). **e**, Schematic view of AAV5-mCherry-*Gfap*-Cre virus injection into spinal cord of CD63-GFP^f/+^ mice and representative images of induced CD63-GFP^+^ and mCherry signals in proximal and distal spinal cord sections from the injection site. Mice analyzed 2 weeks post-injection; Red dashed circles: CD63-GFP^+^ area; White dashed circles: ventral horn grey matter area; mCherry signals are only visible within 500 μm from the injection site. Scale bar: 200 μm; **f**, Quantification of the distance CD63-GFP^+^ signal traveled along spinal cord in AAV5-mCherry-*Gfap*-Cre injected CD63-GFP^f/+^ mice. n = 4 mice (injected at P90); **g**, Representative immunoEM images of hCD63 labeling on spinal cord sections of AAV5-mCherry-*Gfap*-Cre-injected CD63-GFP^f/+^ mice. Intracellular immunogold signals (yellow arrows) are observed inside astroglia (astro., subpanel i) and in neuronal post-synaptic (indicated with black arrows, subpanel iv) dendritic compartment (D, subpanel iv). Subpanels ii and iv are the magnified views of subpanels i and iii, respectively. A: axonal terminal; Mt: mitochondria; Scale bars, 100 nm. p value in **c** determined from two-tailed t test.

To further examine astroglial exosome distribution and spreading in spinal cord, we performed stereotaxic injections of AAV5-mCherry-*Gfap*-Cre virus (0.5μl, 4 x 10^12^ gc/mL) into the grey matter of lumbar spinal cord of adult (P90) CD63-GFP^f/+^ mice. We decided to perform injections on adult mice to better target spinal cord grey matter which is nearly unfeasible in P1 pups. However, we also observed widespread CD63-GFP^+^ puncta from astroglia (Supplementary Fig. 4b i) that surround βIII-tubulin^+^ axons (Supplementary Fig. 4b ii) in longitudinal spinal cord sections of young (P7) *Slc1a3*-CreER^+^CD63-GFP^f/+^ mice following a 4-OHT injection (at P2), suggesting that abundant astroglial exosomes are secreted in the spinal cord during the first postnatal week. We also performed HepaCAM immunostaining on spinal cord sections of P7 4-OHT-injected *Slc1a3*-CreER^+^CD63-GFP^f/+^ mice and observed HepaCAM immunoreactivity co-localized with CD63-GFP^+^ puncta (white arrows, Fig. 4d). HepaCAM protein expression in spinal cord was observed as early as P0 that also undergoes a similar developmental up-regulation (Fig. 4c-d) as in cortex^16^.

A single AAV injection into the spinal cord of adult mice results in bright CD63-GFP^+^ fluorescence at the injection site, indicated by the mCherry fluorescence (yellow arrow, Fig. 4e) expressed from the AAV. By quantifying and calculating the percentage of CD63-GFP^+^ area (red dashed circle) out of the ventral horn grey matter (GM) area (white dashed circle) on coronal sections, we found that CD63-GFP^+^ signals spread as far as 4000 μm in each direction along the spinal cord (Fig. 4f) while the AAV (indicated by mCherry) only diffuses around 500 μm in each direction (Fig. 4e). This longitudinal CD63-GFP^+^ signal analysis from the focal AAV injection suggests that astroglial exosomes are indeed able to spread over long distances. To overcome the detection limit of confocal microscopy, we further examined induced hCD63 signals by immuno-EM in spinal cord sections of AAV5-mCherry-*Gfap*-Cre-injected CD63-GFP^f/+^ mice. Clustered hCD63^+^ immunogold signals were found not only inside astroglia (yellow arrows, labeled ILVs or MVBs, Fig. 4g ii) but also in post-synaptic (indicated by black arrows) dendritic (“D”) terminals (yellow arrows, Fig. 4g iv), further supporting the notion that CD63-GFP^+^ A-Exo. are indeed able to be secreted extracellularly and subsequently be internalized into neurons.

### HepaCAM is important for early postnatal CST axon growth and promotes growth cone size

Although developing axon growth in the CNS is mostly completed at birth in mice, CST axons continue to grow especially during the 1^st^ postnatal week^2^ (Fig. 5a diagram) during which A-Exo. are abundantly secreted (Fig. 4b). Anterograde tracing dyes, such as CM-DiI, have been previously used^37^ to label layer V pyramidal neurons in the motor cortex and their descending axons, especially during early postnatal development (representative labeling image in Fig. 5b), which allows tracing of their continuous postnatal growth. Other genetic approaches, such as Emx1-Cre x Thy1-STOP-YFP^38^ or UCHL1-eGFP^39^ mice, are specifically suitable for adult but not developing CST labeling and can also be non-specific^40^. We therefore performed focal CM-DiI dye injections on the layer V motor cortex of WT and HepaCAM KO pups (P1). Pups were collected 48h post injection and longitudinal sections of the spinal cord were prepared as shown in Supplementary Fig. 5a. This time point was chosen to facilitate the preparation of longitudinal spinal cord sections and to observe consistent DiI labeling. CM-DiI-labeled CST axons undergo pyramidal decussation (PD, orange arrows, Fig. 5c) and continue to elongate in spinal cords. The representative images (Fig. 5c) were created by superimposing multiple individual images taken from longitudinal spinal cord sections from lateral to middle orientation (Supplementary Fig. 5a-b). CST axons cross the midline and begin to enter the spinal cord at birth in mice. We therefore quantified the length (between two yellow lines, Fig. 5c, i-ii) of CM-DiI-labeled CST axons that grow into the spinal cord from the PD. Quantitative measurement found that CST axons grow a significantly (∼1300 μm, p = 0.01) shorter distance into the spinal cord of HepaCAM KO pups when compared to WT mice from P1 to P3 (Fig. 5d). This is consistent with the *in vitro* results that HepaCAM-deficient A-Exo. only modestly promote axon growth compared to HepaCAM-expressing A-Exo. (Fig. 3a). In parallel, a recent study showed that the loss of HepaCAM in astroglia has no effect on density of excitatory intracortical or thalamocortical synapses and only modestly decreases the density of inhibitory synapses in layer I cortex^19^.

**Fig. 5.**
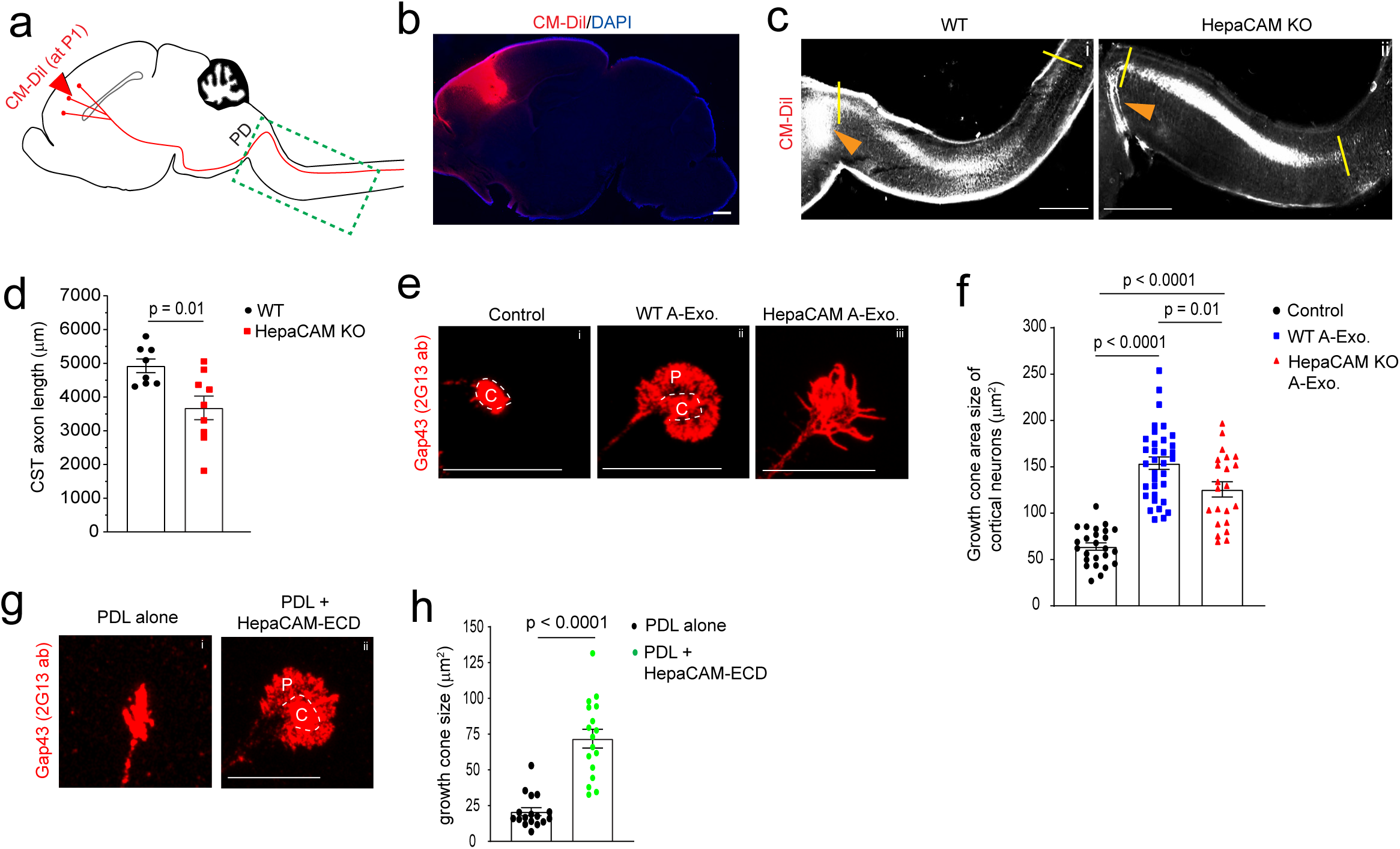
HepaCAM is essential for early postnatal CST axon growth and expands axon growth cone size. **a**, Diagram of CM-DiI dye injections at the motor cortex to label layer V pyramidal neurons and descending CST axons; PD: pyramidal decussation; green dashed box indicates postnatal CST growth (as shown in panel c); **b**, Representative image to show the CM-DiI labeling in the motor cortex 2 days following the injection; Scale bar: 1mm; Representative images (**c**) and quantification (**d**) of CM-DiI-labeled CST axons in the spinal cord of WT (i) and HepaCAM KO (ii) mice. Orange arrows indicate the pyramidal decussation; yellow lines indicate the beginning and ending points for the CST axon length measurement; The image was generated by superimposing images of serial longitudinal sections, which are shown in Supplementary Fig. 5a. Scale bar: 1mm; n = 8-9 mice/group; Representative images (**e**) and quantification (**f**) of axon growth cone size of control (i) cortical neurons or cortical neurons treated with WT (ii) or HepaCAM KO (iii) A-Exo. C: center domain (white circle); P: peripheral domain (growth cone area outside of the center domain); Scale bars: 20 µm. n = 14-44 neurons (3 biological replicates)/group; Representative images (**g**) and quantification of axon growth cone size (**h**) of cortical neurons grown on either PDL alone (i) or PDL/HepaCAM-ECD (ii) coating. n = 17 neurons (3 biological replicates)/group; Scale bar: 20 μm; p value in **d** and **h** determined from two-tailed t test; p values in **f** determined using one-way ANOVA followed by a Tukey post-hoc test.

Axon elongation is primarily driven by the growth cone, which is composed of central and peripheral domains^9^. In particular, the peripheral domain of the axon growth cone is abundant with actin filament-organized filopodia and lamellipodia^9^. It has been well established that actively elongating axons, such as under nerve growth factor (NGF) stimulation, have increased peripheral domain size in growth cones, an indication of extended filopodia and lamellipodia^41^, that increases their contact with surrounding substrates^9^. In contrast, collapsed growth cones have these filopodia and lamellipodia retracted, leading to reduced or even lost peripheral domains^42^. To determine whether A-Exo. alter axon growth cone morphology and especially peripheral domain size, we performed immunostaining of growth associated protein 43 (GAP43, 2G13 clone antibody) that specifically labels axon growth cones^43^ following A-Exo. treatment of cortical neurons. Co-immunostaining of Tau and Map2 confirmed specific axon growth cone labeling revealed by the 2G13 antibody (Supplementary Fig. 5c). As expected, axon growth cones from untreated control neurons (DIV 6) have minimal peripheral domains (Fig. 5e i) following initial growth on PDL/laminin (LN) coated coverslips. A-Exo. treatment induces an enlarged fan-shaped growth cone morphology with extended peripheral domain (“P”, Fig. 5e ii), which is a characteristic growth cone morphology induced by CAM substrates, but not LN substrates, which leads to multiple and long protrusions of filopodia in the peripheral domain^41, 44^. In contrast, growth cone morphology of neurons treated with HepaCAM-deficient A-Exo. tends to have protrusions of filopodia (Fig. 5e iii) and the total growth cone size is also significantly reduced (Fig. 5f). To directly test the effect of HepaCAM on axon growth cone morphology and size, we next examined axon growth cones of cortical neurons cultured on PDL alone (to minimize the influence of the laminin substrate on growth cone) or on PDL/HepaCAM ECD coating. Consistent with the strong stimulation of axon growth by HepaCAM ECD coating (Fig. 3e-f), HepaCAM ECD induces the formation of a large peripheral domain in growth cones (Fig. 5g ii). The overall growth cone size of neurons treated with HepaCAM ECD is 3-fold larger (p < 0.0001) than that of neurons grown on PDL alone (Fig. 5h). The changes in axonal growth cone morphology and size observed after WT, HepaCAM-deficient A-Exo., and HepaCAM ECD treatment support the direct function of HepaCAM in regulating axonal growth cones.

### ApoE in non-exosome ACM fractions inhibits A-Exo. -mediated stimulation on neuronal axon growth

Although HepaCAM on A-Exo. robustly stimulates axon growth, synaptogenesis but not axon growth was primarily observed in neurons stimulated by ACM or co-cultured with astrocytes in previous studies^45^. We also confirmed that non-exosome FT from ACM has no stimulatory effect on axon growth (Fig. 1d). Intrigued by these observations, we decide to test the possibility that non-exosome fractions of ACM may suppress A-Exo’s effect on axon growth. Interestingly, mixing of 0.2x (concentrated from 2mL, 70μg proteins) and 0.5x (concentrated from 5mL, 175ug proteins) non-exosome ACM flow-through (FT) from the SEC column with A-Exo. completely abolishes A-Exo’s stimulatory effect on axon growth (Fig. 6a-b). Subsequent immunoblotting further found that ApoE and ApoJ, two known apolipoproteins secreted from astrocytes, are either mostly (> 98% for ApoE) or completely (ApoJ) detected only in non-exosome fractions of ACM (Fig. 6c) with very low ApoE (but not ApoJ) immunoreactivity detected in A-Exo. only after oversaturated exposure (Supplementary Fig. 6a). Other apolipoproteins, such as ApoB, were not detected in ACM (Fig. 6c). We then mixed human (h)APOE3 with A-Exo. and found that hAPOE3 is able to dose-dependently abolish the stimulatory effect of A-Exo. on axon growth (Fig. 6d-e). The inhibitory dose of hAPOE3 (starting at 10μg/mL) is comparable to the ApoE concentration in ACM (∼15μg/mL) based on the densitometry of ApoE immunoblotting with human APOE and ACM samples. Previously, three major APOE protein isoforms, APOE2, 3, and 4, have been identified that are closely associated with Alzheimer’s disease (AD) risks in human^46^. However, these differential APOE protein isoforms equally and strongly inhibit A-Exo’s stimulatory effect on axon growth (Fig. 6f). In addition, this inhibitory effect is specifically mediated by hAPOE but not by hAPOB or hAPOJ (Supplementary Fig. 6b).

**Fig. 6.**
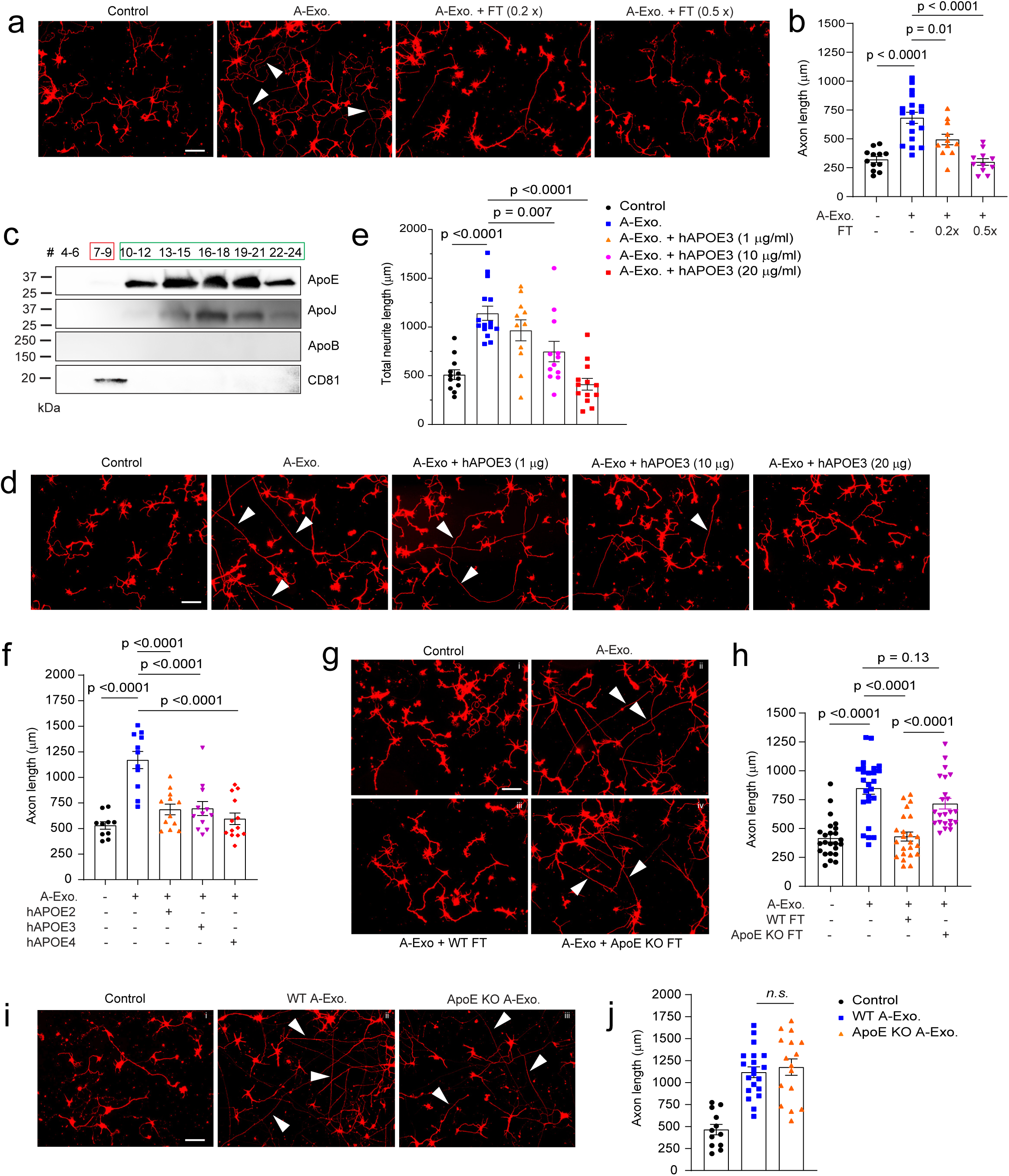
ApoE in non-exosome ACM fractions inhibits A-Exo. -mediated stimulation on neuronal axon growth. Representative images (**a**) and quantification (**b**) of βIII-tubulin^+^ neuronal axon (white arrows) length following treatment of cortical neurons with A-Exo. or A-Exo. mixed with flowthrough (FT) from the SEC column; 0.2x and 0.5x FT each is concentrated from 2- or 5-mL exosome-free ACM, respectively. n = 10-18 neurons (from 2 biological replicates)/group; Scale bar: 100 μm; **c**, Representative immunoblot of different apolipoproteins in all eluted fractions (500 μl/fraction, pooled as indicated) of ACM (100 mL) from SEC with optimal exposure. Unconcentrated elution (15 μl/sample) was run on immunoblot; Representative images (**d**) and quantification (**e**) of βIII-tubulin^+^ neuronal axon (white arrows) length following co-treatment of cortical neurons with A-Exo. and different dose of hAPOE3. n = 10-15 neurons (3 biological replicates)/group; Scale bar: 100 μm; **f**, Quantification of βIII-tubulin^+^ neuronal axon length following co-treatment of A-Exo. with common hAPOE isoforms. n = 11-13 neurons (2 biological replicates)/group; Representative images (**g**) and quantification (**h**) of βIII-tubulin^+^ neuronal axon (white arrows) length in control cortical neurons (i) or neurons treated with A-Exo. (ii) and A-Exo. mixed with WT (iii) or ApoE KO (iv) FT, respectively. Scale bar: 100 μm; n=21-23 neurons (3 biological replicates)/group; Representative images (**i**) and quantification (**j**) of βIII-tubulin^+^ neuronal axon (white arrows) length in control (i) cortical neurons or neurons treated with WT (ii) or ApoE KO (iii) A-Exo. Scale bar: 100 μm; 1μg A-Exo. was used in each treatment. p-values in **b**, **e**, **f, h,** and **j** were calculated using one-way ANOVA followed by a Tukey post-hoc test; *n. s.*: not significant.

Lipids, including cholesterol, are important structural building blocks for developing axon growth^47^. Although lipids are primarily synthesized within neurons (either in cell bodies or locally at axons) and anterogradely transported to axons for developmental growth^47^, during axon regeneration, ApoE, the primary cholesterol carrier, has been shown to contribute to axon growth^48^. We directly added ApoE, cholesterol, and hHDL separately to cultured neurons to test whether they can stimulate axon growth. Interestingly, none of these treatments had any effect in promoting axon growth of cortical neurons (Supplementary Fig. 6c). Additionally, co-treatment of neurons with A-Exo. and receptor associated protein (RAP), a competitive inhibitor for ApoE binding to its receptor low density lipoprotein receptor-related protein 1 (LRP1) for cholesterol delivery, also has no effect on A-Exo’s stimulation of axon growth (Supplementary Fig. 6d-e), suggesting that ApoE/cholesterol does not mediate the stimulatory effect of A-Exo. on axon growth. This is also consistent with the very low level of ApoE detected on A-Exo. (Supplementary Fig. 6a). To confirm that ApoE in ACM indeed inhibits A-Exo’s stimulatory effect on axon growth, we collected ApoE-deficient ACM from ApoE KO mouse pups. The loss of ApoE in ApoE KO ACM and A-Exo. was confirmed by immunoblot (Supplementary Fig. 6f). Consistently, FT from the ApoE KO ACM has no inhibitory effect on A-Exo’s effect on axon growth while wild type (WT) FT completely inhibits A-Exo’s effect on axon growth (Fig. 6g-h). As we showed above that HepaCAM is essential in mediating A-Exo’s axon growth stimulation, we then tested whether ApoE physically binds to HepaCAM to block its interaction with neurons. However, no ApoE was detected in HepaCAM pull-down from astroglial cell lysate, while in the IgG control HepaCAM was not pulled down nor ApoE was detected (Supplementary Fig. 6g), which suggests no direct binding between HepaCAM and ApoE. Meanwhile, ApoE KO A-Exo. stimulate axon growth similarly as WT A-Exo. (Fig. 6i-j), further suggesting that ApoE is not involved in mediating A-Exo’s stimulatory effect on axon growth. Taken together, these results demonstrate that ApoE is minimally found in A-Exo. and not involved in A-Exo’s stimulatory effect on axon growth; rather, ApoE is highly abundant in the non-exosome ACM fraction that strongly inhibits A-Exo’s stimulatory effect on axon growth.

### ApoE deficiency reduces developmental synaptogenesis and dendritic spine formation on cortical pyramidal neurons *in vitro* and *in vivo*

ApoE-mediated transport of cholesterol to neurons has been shown to promote synaptogenesis in cultured RGCs ^4^. To examine whether this ApoE/cholesterol pathway is also essential for cortical neuronal synaptogenesis, WT cortical neurons were treated with ACM collected from WT or ApoE KO astrocyte cultures. The loss of ApoE leads to accumulated cholesterol in cultured ApoE KO astrocytes, indicated by Filipin 3 staining (Supplementary Fig. 7a iii, Supplementary Fig. 7b), similar to the results of treatment with U18666A, an inhibitor for cholesterol transport, in WT astrocyte cultures (Supplementary Fig. 7a ii), suggesting a reduced secretion of cholesterol from ApoE-deficient astrocytes. Consequently, cortical neurons treated with WT ACM have strongly increased VGluT1 and PSD95 puncta density on the neurites (Fig. 7a ii, Fig. 7b-c). However, only a modest increase in VGluT1 (p = 0.05) and PSD95 (p = 0.14) puncta density was observed in neurites of cortical neurons treated with ApoE-deficient ACM, suggesting that the astroglial ApoE/cholesterol pathway similarly promotes synaptogenesis of cortical neurons.

**Fig. 7.**
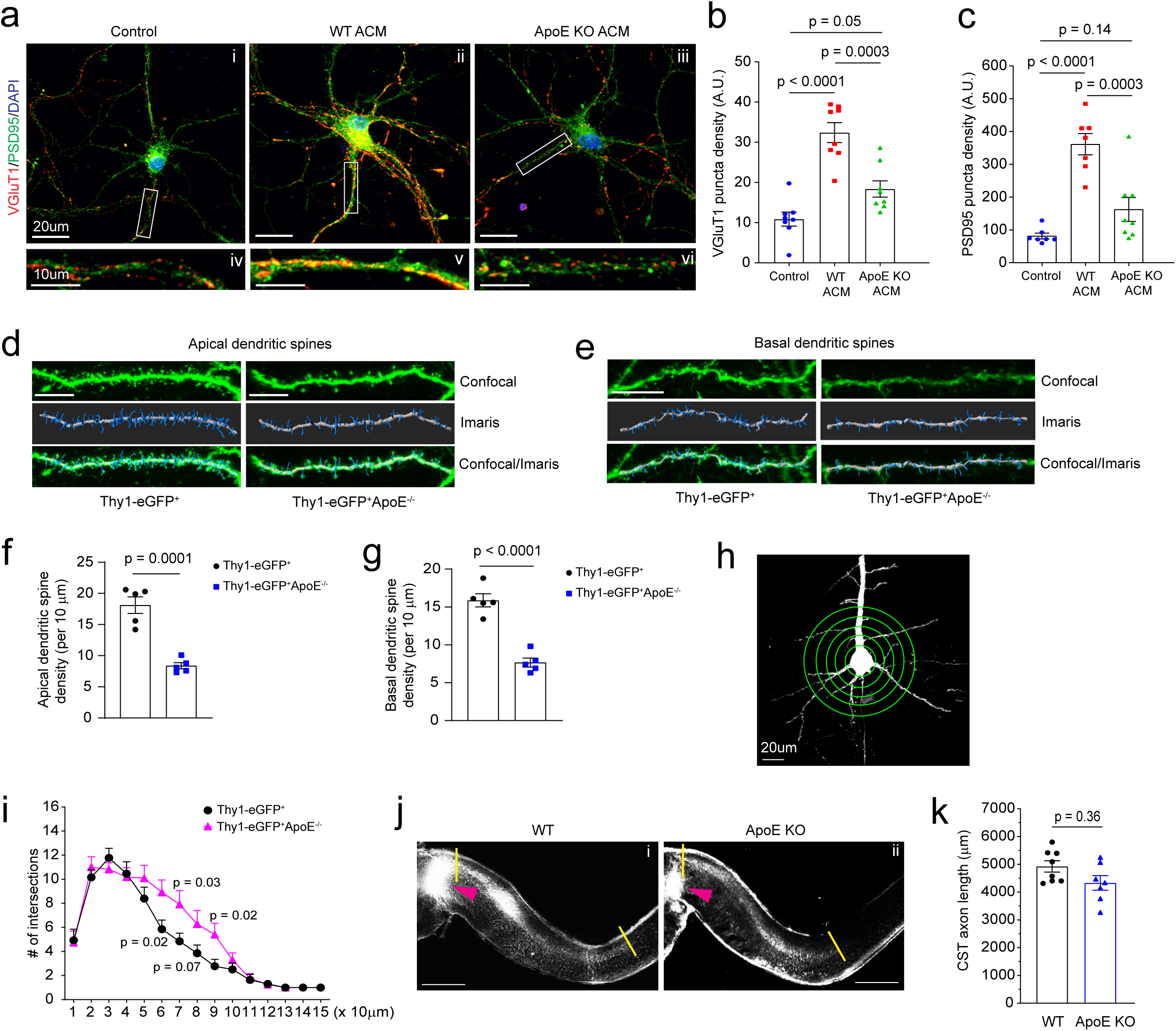
ApoE deficiency reduces developmental dendritic spine formation and alters dendritic branching on layer V cortical pyramidal neurons. Representative image of VGluT1 and PSD95 staining in cortical neuronal cultures (**a**) and quantification of VGluT1 (**b**) and PSD95 density (**c**) on neurites following ACM treatment. Subpanels: control cortical neurons (i) and neurite (iv), cortical neurons (ii) and dendrite (v) treated with WT ACM, and cortical neurons (iii) and dendrite (vi) treated with ApoE KO ACM; n = 7-9 neurons (2 biological replicates)/group; Representative confocal and Imaris images of apical (**d**) and basal (**e**) dendrites and spines of layer V pyramidal neurons from motor cortex of Thy1-eGFP^+^ and Thy1-eGFP^+^ApoE^-/-^ mice (P30). Dendrites and spines were traced and quantified in Imaris. Scale bars: 10 μm; Quantification of apical (**f**) and basal (**g**) dendrites of layer V pyramidal neurons from motor cortex of Thy1-eGFP^+^ and Thy1-eGFP^+^ApoE^-/-^ mice (P30). n = 5 mice/group; Representative neuron image (**h**) and 3D Sholl analysis (**i**) of layer V pyramidal neurons from motor cortex of Thy1-eGFP^+^ and Thy1-eGFP^+^ApoE^-/-^ mice. Scale bar: 20 μm; n = 5 mice/group; Representative images (**j**) and quantification (**k**) of CM-DiI-labeled CST axons in the spinal cord of WT (i) and ApoE KO (ii) mice. Orange arrows indicate the pyramidal decussation; yellow lines indicate the beginning and ending points for the CST axon length measurement; Scale bar: 1mm; n = 8-9 mice/group; p value in **f**, **g**, and **k** determined by two-tailed t test; p values in **b** and **c** determined using the one-way ANOVA followed by a Tukey post-hoc test; p values in **i** determined using the multiple t-test;

ApoE is known to be mostly expressed in and secreted from astroglia in the homeostatic CNS and ApoE mRNA was found to be strongly up-regulated in astroglia during postnatal development by single cell sequencing^49^. We performed ApoE immunoblotting and also found that ApoE protein is only lowly expressed at birth and is robustly up-regulated in cortical tissues during postnatal development during which synaptogenesis occurs (Supplementary Fig. 7c-d). To directly examine whether ApoE deficiency affects dendritic branching and developmental dendritic spine formation of cortical neurons especially layer V pyramidal neurons in the motor cortex, we generated Thy1-eGFP^+^ApoE^-/-^ mice and analyzed pyramidal neuronal morphology and their dendritic spine density between Thy1-eGFP^+^ and Thy1-eGFP^+^ApoE^-/-^ mice. Thy1-eGFP (H line) mice have been widely used to illustrate neuronal morphology including dendritic spines^50^. We observed well-labeled neurons across the CNS including pyramidal neurons in layer V motor cortex (Supplementary Fig. 7e i-ii). The clear eGFP labeling also facilitates clear identification of apical and basal dendrites and spines (Fig. 7d-e). By using the filament tracing function in Imaris software, representative eGFP^+^ dendritic spines from individual layer V pyramidal neurons (Supplementary Fig. 7f) in both Thy1-eGFP^+^ and Thy1-eGFP^+^ApoE^-/-^ mice were traced and quantified. Consistent with our *in vitro* results (Fig. 7a-c), ApoE deficiency leads to substantially reduced spine density on both apical and basal dendrites (Fig. 7f-g) of layer V pyramidal neurons in Thy1-eGFP^+^ApoE^-/-^ mice. Unexpectedly, the loss of ApoE also increased secondary dendritic branches in Thy1-eGFP^+^ApoE^-/-^ mice, based on 3D Sholl analysis (Fig. 7h-i), likely compensating for the reduced dendritic spine density. We further performed CM-DiI injections on ApoE KO pups (P1) to examine whether the loss of ApoE also affects postnatal axon growth, as part of CST extension to the spinal cord, of layer V pyramidal neurons in the motor cortex. We only observed modestly reduced CST axon growth (average ∼600 μm shorter, Fig. 7j) but not statistically significant (p = 0.37, Fig. 7k) in ApoE KO pups compared to WT pups. This is consistent with our finding of no obvious changes of HepaCAM protein expression in ApoE KO mouse cortex, nor no ApoE protein expression changes in HepaCAM KO mice (Supplementary Fig. 7g).

## Discussion

In our current study, by employing an optimized SEC-based exosome isolation procedure, we defined a previously unknown astroglial exosome-dependent regulatory pathway that stimulates developmental pyramidal neuronal axon growth. This pathway is specifically mediated by astroglial exosomes, as exosome-depleted ACM fractions have no effect in stimulating axon growth. The stimulating effect is axon-specific with a primary action on axon growth cones but not affecting dendritic arborization, length, and synaptogenesis. Consistently, SEC-isolated astroglial exosomes are minimally associated with known astroglia-derived soluble proteins that regulate synaptogenesis. This further supports the notion that astroglial exosomes represent a distinct and unique class of secreted signals from astroglia, in contrast to astroglial secretion of soluble proteins and small molecules to modulate synaptogenesis/maturation and transmission ^3^.

Although trophic factors such as NGF/BDNF, are well established to potently promote axon growth ^47^, our proteomic analysis found no trophic factors in astroglial exosomes, ruling out their involvement in astroglial exosome-stimulated axon growth. Our results also showed that neither RNA mechanisms nor endocytosis are involved in the axon growth-stimulating effect of astroglial exosomes, which is distinct from previous reports that miRNA signals can mediate the axon growth-stimulating effect of mesenchymal stem cell (MSC) exosomes or regulate dendrite complexity through endocytosis^25, 51^. Instead, our results provided evidence for an essential and sufficient role of surface HepaCAM on astroglial exosomes in promoting axon growth, representing a unique surface contact mechanism for exosome action. This also provides a mechanism for plasma membrane surface proteins to be secreted through the MVB pathway, as initially observed with the secretion of transferrin receptors in reticulocytes^52^. These prior studies and our results indicate a growing understanding of the diverse mechanisms and effects of cell-type specific exosomes. Our results also revealed an important new function of HepaCAM to mediate intercellular signaling between astroglia and neuronal axons, in addition to its intracellular role as a binding partner to facilitate proper targeting of anion and chloride channels on glial cell surface ^17, 34^ and regulate boundary of neighboring astroglia ^19^. How HepaCAM activates downstream pathways in neurons to expand the surface area of growth cones and to promote axon growth remains unclear. CAM protein-mediated downstream signaling is highly diverse and complex by either activating receptors such as integrins, FGF receptors or directly binding intercellularly^53^. As HepaCAM ECD is sufficient to stimulate axon growth, it is possible that HepaCAM ECD activates its neuronal receptor, which remains to be identified, for downstream signaling. In axon growth cones, anterograde polymerization of actin filaments (F-actins) contributes to retrograde flow of F-actin and pushes the growth cone in the forward direction ^9^. Previous studies have identified several kinases, particularly focal adhesion kinase (FAK), that are activated downstream of CAM proteins, to promote actin polymerization and axon growth^54, 55^. Whether these pathways are involved in HepaCAM ECD’s axon-stimulating effect will be investigated in future studies.

Although astroglia are able to secrete various EVs, these previous studies were almost exclusively carried out in cultures ^25, 26, 28^. By employing our previously generated cell-type specific exosome reporter mice and Ai14 reporter mice, our results illustrated the *in situ* localization and dynamics of secreted A-Exo. in the motor cortex during development and in adult spinal cord. Our results showed that astroglial exosomes are able to spread long distances (up to 8000 μm bidirectionally). In particular, our results showed that A-Exo. can be abundantly localized outside of astroglia during the 1^st^ postnatal week when astroglial morphology remains primitive with limited processes. These extracellularly localized A-Exo. may serve as an alternative cell to cell contact mechanism, especially in the 1^st^ postnatal week, to allow long-range spreading of surface contact signals, such as HepaCAM, via A-Exo. Thus, surface expressed HepaCAM on A-Exo. (Fig. 8) is a mobile astroglial CAM signal to stimulate CST axon growth postnatally. As many synapses (both excitatory and inhibitory) are not ensheathed by astroglial processes even in the adult CNS^3, 56^, mobile surface contact signals on A-Exo. may mediate specific intercellular signaling, in addition to direct plasma membrane contact or the cleavage of transmembrane protein signals.

**Fig. 8.**
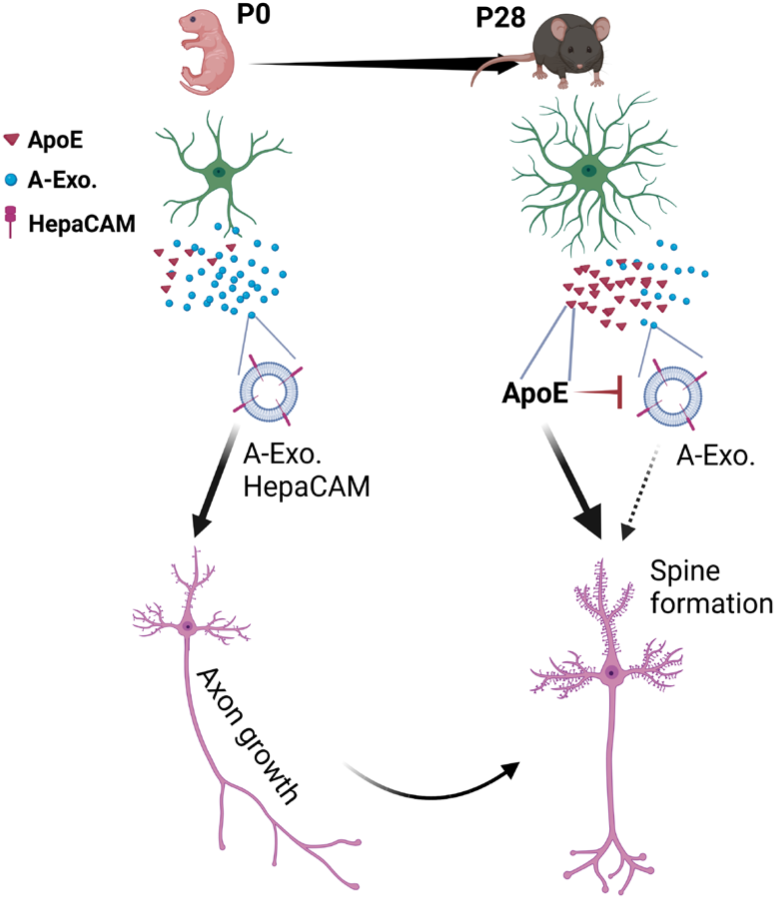
Developmental astroglial exosome HepaCAM signaling and ApoE coordinates postnatal cortical pyramidal neuronal axon growth and dendritic spine formation Abundantly secreted astroglial exosomes promote CST axon growth during early postnatal development (within 1^st^ postnatal week) when ApoE is lowly expressed; this effect is antagonized by increased ApoE expression to promote dendritic spine formation after CST axon growth is completed later during the postnatal development.

ApoE is the major carrier for transporting cholesterol and phospholipids in the CNS. It has been extensively studied in CNS pathology, and human APOE polymorphism has been closely associated with AD pathogenesis ^46^. However, the developmental role of ApoE has not been examined *in vivo*, despite an early study suggesting that ApoE-mediated transport of cholesterol promotes synaptogenesis in cultures ^4^. Our results from both genetic and pharmacological approaches showed that the ApoE/cholesterol pathway is not involved in mediating A-Exo’s stimulatory effect on axon growth. On the contrary, abundant ApoE levels are only found in non-exosome ACM fractions that strongly inhibit the stimulatory effect of A-Exo. on axon growth. Interestingly, our and others’ results showed that ApoE is expressed at low levels during the 1^st^ postnatal week and is highly up-regulated later in development ^49^ which could strongly inhibit A-Exo’s stimulatory effect on axon growth and promote cholesterol transport and subsequent synaptogenesis in cortical pyramidal neurons. Consistently, early postnatal CST axon growth from cortical pyramidal neurons ends around P10 in mice. Thus, these results suggest that ApoE-mediated inhibition of A-Exo’s stimulation on axon growth facilitates the developmental transition of layer V pyramidal neurons in the motor cortex from axon growth to dendritic spine formation (Fig. 8). In support of this notion, we found that ApoE deficiency leads to significantly reduced spine density on both apical and basal dendrites of layer V pyramidal neurons in the motor cortex of ApoE KO mice. These results provide important new insights about the function of ApoE during postnatal CNS development.

How ApoE inhibits the stimulatory effect of A-Exo. on axon growth remains unclear. Although HepaCAM is essential to mediate A-Exo’s stimulation on axon growth, we found no evidence that ApoE binds to HepaCAM to block its stimulation on axon growth. In addition, ApoE can be readily separated from A-Exo. using the SEC with simple PBS wash, also suggesting a non-covalent nature in the interaction between ApoE and A-Exo. Since ApoE has high affinity to cholesterol and phospholipids and these lipids are well distributed on A-Exo. surface, it is conceivable that ApoE interacts with such lipids to block A-Exo’s surface contact with neurons especially growth cones, which will be tested in future studies.

## Supporting information

A-Exo. treated

control (untreated)

## Acknowledgments

We thank Dr. Peter Juo (Tufts University School of Medicine), Dr. Fen-Biao Gao (University of Massachusetts Chan School of medicine), and Dr. Zhigang He (Boston Children’s Hospital) for constructive discussions. We thank Dr. Cagla Eroglu and Dr. Katherine Baldwin (Department of Cell Biology, Duke University School of Medicine) for providing HepaCAM floxed mice. This work was supported by NIH grants RF1AG057882, RF1AG059610, R01NS118747, R01NS125490, and R01AG078728 (YY). Imaging was performed with the assistance of the Tufts Center for Neuroscience Research. EM was performed with the assistance of the Harvard Medical School EM Core Facility. LC/MS/MS and proteomic analysis was performed with the help of University of Massachusetts Medical School.

## Author contributions

SJ designed and performed majority of experiments in this study and wrote the manuscript. YT performed spinal cord injections and CD63-GFP image analysis. XC performed immunostaining, image analysis, and wrote the manuscript. RJ and VP performed image analysis, helped with exosome isolation, and wrote the manuscript. YY designed overall study, analyzed data, and wrote the manuscript.

## Data availability statement

All data supporting this study are available upon request

## Materials and Methods

### Reagents and neuronal culture treatments

Dynasore (Sigma Aldrich), RNase A (Roche), Proteinase K (Fisher Scientific), CM-DiI (Thermo Fisher Scientific, C7001), and human HepaCAM protein extracellular domain (ECD, amino acid sequence 1-240, 16047-H08H) (Sino Biological Inc.) were used in this study. Dynasore (stock 50 mM) was prepared in DMSO and diluted 1000x in neuronal growth medium for treatment. Antibodies (final concentration 100 μg/mL) were mixed with neuronal growth medium and added onto primary neuronal cultures 2 hours before A-Exo. treatment with exosomes. HepaCAM ECD coating is described below. Neuronal treatment with various drugs and/or exosomes was generally at DIV 3-4 for 24h unless specifically described in main text.

### Mice

CD63-GFP floxed mice were generated in the lab by homologous recombination, as previously described ^35^. The WT (C57B/6J, #000664), Ai14-tdT^f/f^ reporter (#007914), ApoE-KO (B6.129P2-Apoe tm1Unc/J #002052), B6.Cg-Tg (Thy1-YFP)HJrs/J (#003782), and B6.C-Tg(CMV-Cre)1Cgn/J(#006054) mice were obtained from the Jackson Laboratory. HepaCAM knock-out (KO) mice were generated by breeding HepaCAM floxed mice (a kind gift from Dr. Cagla Eroglu at Duke University)^19^ with CMV-Cre mice. Both male and female mice were used in all experiments. All mice were maintained on a 12 h light/dark cycle with food and water ad libitum. Care and treatment of animals in all procedures strictly followed the NIH Guide for the Care and Use of Laboratory Animals and the Guidelines for the Use of Animals in Neuroscience Research. Animal protocols used in this study have been approved by the Tufts University IACUC.

### Primary cortical astrocyte and neuronal culture

For cortical astrocyte cultures, P0-P3 mouse pups were decapitated, and cerebral cortices were removed and transferred into astrocyte growth medium (Dulbecco Minimum Essential Medium, DMEM, supplemented with 10% exosome-depleted FBS (fetal bovine serum, Gibco) and 1% penicillin/streptomycin) for dissection on ice. Meninges were stripped and cortices were minced and placed into 0.05% trypsin-EDTA solution for 10 min in a 37 °C water bath. The enzymatic reaction was stopped by addition of astrocyte culture medium. The tissue was washed twice with astrocyte medium and then gently dissociated by trituration with a fire-polished Pasteur pipette. Dissociated cells were filtered through a 70 μm strainer to collect a clear astrocyte cell suspension. Primary cortical neuron cultures were prepared from embryonic day 14-16 mouse brains. In brief, cortices were dissected and dissociated using 0.05% trypsin-EDTA solution for 10 min at 37 °C. Cells were seeded (1∼2 x 10^4^/well) on Poly-D-lysine coated coverslips (Neuvitro), Poly-D-lysine and laminin coated coverslips (GG-12-Laminin Neuvitro) in 24well culture dish (1∼ 2 x10^4^ cells/well) with 1mL neuron plating medium containing DMEM, 10% FBS and 1% Pen-Strep at 37 °C in a humidified chamber of 95% air and 5% CO2. After a 12h seeding period, neuron plating medium was replaced by 700 μL neuron culture medium composed of neurobasal medium (Invitrogen), 2% B27 supplement (Thermo Fisher Scientific), 1% 100x GlutaMAX (Thermo Fisher Scientific), and 1% penicillin–streptomycin (Thermo Fisher Scientific). As we only observe < 5% astrocytes in neuronal cultures and neuronal cultures were collected by DIV 7-8 at the latest, mitotic inhibitors such as cytosine arabinoside (ara-C) were not used in neuronal cultures.

### Intracellular cholesterol staining and quantification in primary astrocyte cultures

Intracellular cholesterol levels were measured with the cell-based cholesterol assay kit (Abcam, ab133116) Briefly, primary culture astrocytes were fixed with 4% PFA for 10 min. Astrocytes were stained with Filipin 3 according to manufacturer instructions. GFAP (rabbit anti-GFAP, 1:1000, Dako) immunostaining was also performed in primary astrocyte cultures (secondary antibody: anti-rabbit Alexa Fluor 555). Both GFAP immunostaining and Filipin 3 (with DAPI filter) signals were captured with the Zeiss Axio microscope (Zeiss, Heidelberg, Germany) using a 20X objective lens. Filipin 3 signals within individual GFAP^+^ astrocyte that is outlined in ImageJ–FIJI was measured.

### Stereotaxic injections of AAV

For mouse (P30) spinal cord injections, AAV5-mCherry-*Gfap*-Cre virus was obtained from the University of North Carolina Vector Core (Chapel Hill, NC). Spinal cord ventral horn injections were performed with a Hamilton Neuros Syringe with 33G, point style 4, 45-degree bevel needle on a stereotaxic apparatus (Stoelting). A single dose of AAV5-mCherry-*Gfap*-Cre (0.5 μL, 4 x 10^12^ genome copy (gc)/ mL) was injected into the L1 segment of CD63-GFP^f/+^ mice, posterior to the median sulcus 0.4mm laterally, 1.4 mm deep. Injections were performed at a rate of 0.1μL/min. Post-operative care included injections of buprenorphine according to the IACUC requirement. For mouse pups (P1) motor cortex injections, CD63-GFP^f/+^Ai14-tdT^f/+^ mouse pups were anaesthetized on ice for 3 min and then placed in a stereotaxic frame. AAV5-mCherry-G*fap*-Cre (0.3μL, 4 x 10^12^ gc/mL) was stereotaxically injected into the right side of motor cortex (x = 1.0 mm, y=1.8mm, z = 0.6 mm) using a 33-gauge needle. Injections were performed at a rate of 0.1mL/min.

### Exosome purification and qNano particle analysis

Exosomes were prepared from astrocyte conditioned medium (ACM) from primary astrocyte culture (initial seeding: 4×10^6^ cells/10 cm dish). After astrocytes become > 90% confluent, the normal astrocyte growth medium was replaced with exosome depleted astrocyte growth medium composed of DMEM, 10% exosome-depleted FBS (Thermo Fisher Scientific), and 1% penicillin/streptomycin. ACM was replaced and collected every 3 days for up to 4 times (10 mL/10 cm dish). ACM was first spun at 300 x g for 10 minutes at room temperature to remove suspension cells, then at 2,000 x g for 10 minutes at 4°C to remove cell debris, then underwent following purification steps or stored at -80°C. For ultracentrifugation (UC)-based purification, ACM was centrifuged at 10,000 x g for 60 minutes at 4°C. The supernatant was passed through a 0.22 μm polyether sulfone (PES) filter (Merck Millipore, MA, USA) followed by ultracentrifugation at 100,000 x g for 60 minutes at 4°C (SW 41 Ti Rotor, Beckman Coulter Inc). For size-exclusion chromatography (SEC) based isolation, ACM supernatant was first concentrated (to 500 μl) by centrifugation at 3,500 x g for 30 min at 4°C using Centricon® Plus-70 Centrifugal Filter Devices with a 10k molecular weight cutoff (MilliporeSigma). Then the concentrated supernatant was passed through a 0.22 μm PES filter. The qEV original 35nm columns (Izon Science, MA, USA) were then used according to the instructions of the manufacturer. Briefly, the column was rinsed with filtered PBS, and then 500 μl of concentrated and filtered supernatant from ACM was layered onto the top and each eluted fraction (500 μl/fraction) was collected. The eluted fractions were combined, as indicated in text and figure legend, and further concentrated using the Amicon Ultra-4 Centrifugal Filter Units (MilliporeSigma) in certain experiments. Tunable resistive pulse sensing (TRPS) by qNano particle analyzer (Izon Science, MA, USA) was used to measure the size distribution and quantity of isolated exosomes. 15 μl of concentrated and filtered ACM (500 μl from 10 mL/sample) or calibration particles included in the reagent kit were placed in the Nanopore (NP150, Izon Science). Samples were measured at 44∼45 mm stretch with a voltage of 0.6∼0.8 V at 1-pressure levels of 10 mbar. Particles were detected in short pulses of the current (blockades). The calibration particles were measured directly after the experimental sample under identical conditions. The data was processed using the Izon software (version 3.2).

### Exosome and HepaCAM coating for neuronal cultures

Sterile PDL or PDL/LN coated coverslips (Neuvitro) were rinsed twice with 1xPBS, then astroglial exosomes (1 μg) purified from 10 mL ACM were evenly added onto the top of coverslips and incubated for 1 hour in a 37°C cell culture incubator. Coverslips were then washed twice with 1x PBS before use. To block HepaCAM on exosome surface, 100 μg/mL of HepaCAM antibody (ProteinTech) was added separately on top of exosome coated coverslips and incubated for 1 hour at 37°C, then washed twice with 1x PBS before use. For HepaCAM-ECD coating, HepaCAM-ECD protein was diluted with PDL solution to 50 μg/mL, then 80μl PDL/HepaCAM-ECD solution was added onto sterilized coverslips and incubated for 2 hours at room temperature. Coverslips were then washed twice with sterilized water before use.

### Biochemical treatment of exosomes

1 μg A-Exo. (50 μl) was used in each treatment. For RNase treatment, RNase (Roche) was added to A-Exo. at a final concentration of 10 μg/mL for 5 minutes at 37°C, then 20U SUPERase-In RNase inhibitor (Invitrogen) was added to block RNase activity. For proteinase K treatment, proteinase K was added to A-Exo at a final concentration of 10 μg/mL for 5 min at 37°C, then 1% proteinase inhibitor cocktail (P8340, Sigma-Aldrich) was added. For treatment involving sonication, A-Exo. were sonicated at 50 Hz for 30 seconds on ice before RNase or proteinase K treatment. The reaction was washed 2 times using Amicon Ultra Centrifugal Filters (30K MWCO, EMD Millipore) with 1x PBS to remove lysates. A final volume of 60 μl A-Exo. for the various treatments was then added to the primary neuronal culture.

### Immunocytochemistry, immunohistochemistry, live-cell, and confocal imaging

For immunocytochemistry, cultured neurons were fixed in 4% paraformaldehyde for 15 min and permeabilized with 0.2% Triton X-100 for 5 min. The cells were blocked in 3% bovine serum albumin for 30 min and incubated with the following primary antibodies overnight at 4°C: β-III tubulin (1:1000, MAB1195, R&D system), rabbit anti-MAP2 (1:1000, GeneTex), Gap43 Antibody (1:500, Novus Biologicals, clone 2G13), anti-mouse Tau (1:500, GeneTex), mouse anti-Map2 (1:1000, Sigma, M9942), rat anti-GFAP (1:5000, zymed, 273756), rabbit anti-GFAP (1:1000, Dako), and anti-human Tau (1:500, Dako). After incubation with the primary antibodies, neurons were washed three times with PBS, and incubated with following secondary antibodies for 1 h at room temperature: anti-mouse Alexa Fluor 488, anti-rabbit Alexa Fluor 568 and anti-goat 647 Alexa Fluor (1:1000, Invitrogen), and mounted with Prolong™ Glass Antifade Mountant with NucBlue™ Stain (Invitrogen). For immunohistochemistry, mice were anaesthetized with a ketamine/xylazine cocktail and perfused with ice-cold PBS followed by ice-cold 4% paraformaldehyde. Dissected brains were post-fixed overnight in 4% paraformaldehyde at 4 °C for 24 hours, and cryoprotected in 30% sucrose until tissue sinks. The tissue was embedded in OCT compound (Tissue-Tek) and 20 μm tissue sections were cut with a cryostat (Microm HM525). The following antibodies were used: GFAP (1:5000, Dako, #Z0334) and Hepacam (1:200, R&D, #MAB4108). Primary antibodies were visualized with appropriate secondary antibodies conjugated with Alexa fluorophores (1:1000 Invitrogen) and mounted with Prolong™ Gold Antifade Mountant with DAPI (Invitrogen). Low magnification images were taken using the Zeiss Axio Imager fluorescence microscope, using the ZEN2 software to acquire and process the images. Confocal images were taken using the Leica SP8 FALCON confocal laser scanning microscope (15-20 μm Z stack with 0.5 μm step) magnified with 63X (numerical aperture 1.0) objectives; images were processed with LAS X software. Live-cell imaging of primary cortical neurons was performed on a Leica SP8 microscope 24h following the addition of astroglial exosomes (1 μg). The microscope was equipped with a stage top incubator (model: INUBG2A-GSI2X TOKAI HIT) with temperature and CO_2_ control to maintain an environment of 37 °C and 5% CO_2_. The images were taken with a 10x objective len every 3 min for 8 hours using the same exposure time.

### Immunoblotting and immunoprecipitation

Mouse spinal cord, primary astrocyte pellets, and exosome fractions were homogenized with lysis buffer (Tris-HCL pH 7.4, 20 mM, NaCl 140 mM, EDTA 1 mM, SDS 0.1%, Triton-X 1%, Glycerol 10%). Protease inhibitor cocktail (P8340, Sigma) and phosphatase inhibitor cocktail 3 (P0044, Sigma) was added in a 1/100 dilution to lysis buffer prior to tissue homogenization. Total protein amount was determined by DC™ Protein Assay Kit II (Bio-Rad), then lysates were loaded on 4-15% Mini-PROTEAN TGX Stain-Free Protein Gels (Bio-Rad). Separated proteins were transferred onto a PVDF membrane (Bio-Rad) with the Trans-Blot Turbo Transfer System (Bio-Rad). The membrane was blocked with 5% fat-free skim milk in TBST (Tris buffer saline with 0.05% Tween 20) or SUPERBLOCK T20 (TBS) Blocking Buffer (Thermo Fisher Scientific) then incubated with appropriate primary antibody overnight at 4°C. The following primary antibodies were used: Thrombospondin (TSP)-1 (1:100, Santa Cruz clone SC-8), Thrombospondin (TSP)-2 (1:100, BD Biosciences), Hevin (1:200, R&D Systems), Sparc (1:200, R&D Systems), Sema3a (1:200, clone A-18 Santa Cruz), TSG101 (1:100, clone C-2 Santa Cruz), GFAP (1:2000, Dako, #Z0334), mouse CD63 (1:200, MBL, # D263-3), CD81(1:1000, clone B-11, Santa Cruz), β-actin (1:1000, A1978, Sigma), HepaCAM (1:500, ProteinTech), ApoE (1:1000, ABclonal, #A16344), ApoB (1:500, ABclonal, #A4184), ApoJ (1:500, ABclonal, #A1472). Secondary antibodies, including ECL anti-mouse IgG (1:10000, GE HealthCare NA931V), anti-rabbit IgG-HRP (1:5000, GE Health Care NA934V), mouse anti-Goat IgG-HRP (1:1000, Santa Cruz) and anti-Rat IgG-HRP (1:5000, Thermo Fisher Scientific SC-2357) were diluted with Super Blocking Buffer. Bands were visualized on ChemiDoc MP imaging system (Bio-Rad) with ECL Plus chemiluminescent substrate (Thermo Fisher Scientific) or Clarity Max Western ECL Substrate (Bio-Rad).

For immunoprecipitation, Dynabeads® M-270 Epoxy beads (Thermo Fisher Scientific) with anti-CD81 (clone Eat-2, BioLegend), anti-HepaCAM (Affinity Biosciences, # DF12075), and mouse IgG1 (clone MG1-45 BioLegend) was conjugated individually according to the instructions of the Dynabeads Antibody Coupling Kit (Thermo Fisher Scientific). Dynabeads (0.5mg) were mixed with each antibody (5-10μg) and incubated overnight at 4°C with gentle agitation. Beads were then washed with washing buffer and 1xPBS. Concentrated ACM (500μl, from 20 mL/sample) or exosomes isolated from SEC (2 μg/sample) were added and incubated overnight at 4°C with rotating. IP mixes were then placed on a magnetic rack, washed 3 times, and eluted with western blot lysis buffer.

### LC-MS/MS proteomics and data analysis

Three biological exosome samples (20 μg/sample) were separated on 4-15% mini-protein TGX precast protein gels (Bio-Rad) and subsequently stained with Coomassie Blue, then each sample lane was excised and digested with trypsin and spiked with 0.2 pmol of ADH peptides (YEAST Alcohol dehydrogenase 1) at the Mass Spectrometry Facility at the University of Massachusetts Medical School. The samples were then injected into Orbitrap Fusion Lumos Mass Spectrometer (Thermo Fisher Scientific) in technical triplicates for label-free quantitation (LFQ) analysis. The data was searched against Swiss-Prot Mouse protein database using Mascot search engine through Proteome Discoverer software. The data was exported and normalized as intensity-based absolute quantification (iBAQ) quantitative values in Scaffold (version Scaffold_4.10, Proteome software). The selected parameters for protein identification were the following: Protein Threshold > 95%; minimum 3 peptides per candidate protein; Peptide Threshold > 90%; > 1 x 10^5^ iBAQ value in at least one of samples. The iBAQ value of the housekeeping protein ADH was used for normalization of biological replicates.

### Immuno-electron microscope (EM) imaging

EM imaging was performed in the Harvard Medical School Electron Microscopy Facility. AAV5-mCherry-*Gfap*-Cre injected CD63-GFP^f/+^ mice were perfused with 4% paraformaldehyde (PFA) and 0.1% glutaraldehyde. The spinal cord tissue was dissected out and post-fixed in 4% PFA for overnight, then spinal cord slices (100-200 μm) were prepared using vibratome and floated in PBS + 0.02M glycine for 15 minutes. The slices were quenched, permeabilized, and blocked with blocking buffer (1% bovine serum albumin, 0.1% Triton-X100) at 4 °C. Anti-human CD63 (BD Pharmingen, #556019) antibody was then added and incubated overnight at 4°C. The slices were washed three times for 20 min in PBS. The slices were incubated with Protein A-gold 5 nm (1:50, Utrecht, the Netherlands) for 1 hour at 25°C, washed in PBS and fixed in 1% (v/v) glutaraldehyde in PBS for 30 min. For Epon embedding, slices were incubated in 0.5% (w/v) osmium in ddH2O for 30 min, washed three times in ddH2O and then stepwise dehydrated (each step for 10 min) in 70% (v/v) ethanol, 95% (v/v) ethanol, and two times in 100% (v/v) ethanol. The slices were incubated in propyleneoxide, infiltrated in 50/50 propylenoxide/TAAB Epon, embedded in fresh TAAB Epon (Marivac Canada Inc) and polymerized at 60°C for 48h. The block was cut into 60 nm ultrathin sections using a Reichert Ultracut-S microtome. The slices were picked up on to copper grids that have been stained with uranyl acetate and lead citrate. Samples were examined using a JEOL 1200EX transmission electron microscope. Images were recorded with an AMT 2k CCD camera at 30000x magnification. For eluted fractions from SEC columns, negative staining was performed. Briefly, 5µl of the sample was adsorbed to a carbon coated grid that had been made hydrophilic. The primary antibody used was anti-human CD63 (1:20, BD Pharmingen 556019), then samples were incubated with rabbit anti rat bridging antibody (1:50, Abcam ab6703) and Protein A-gold 10nm (University Medical Center Utrecht, the Netherlands). Excess liquid was removed with filter paper (Whatman #1) and the samples were stained with 1% uranyl acetate. The grids were examined in a JEOL 1200EX transmission electron microscope and images were recorded with an AMT 2k CCD camera.

### Image analysis

For neurite tracing and Sholl analysis, neurites and axons were traced and then measured using the Simple Neurite Tracer (SNT) plugin in Fiji ImageJ. Axons were defined as β-III tubulin^+^Map2^-^ neurites. Thy1-YFP^+^ pyramidal neurons in the layer V of motor cortex were traced with SNT in ImageJ. *In vivo* 3D neuronal Sholl analysis was performed on basal dendrites, the radius increment was set at 10 μm. Axon growth cone size was determined in Fiji ImageJ by manually tracing and measuring the area of regions of interest (ROIs) based on the anti-GAP43 antibody fluorescence at the tip of β-III tubulin^+^ (or Tau^+^) Map2^-^ axons. For quantification of VGluT1 or PSD95 puncta, confocal images were taken using the Leica SPE confocal laser scanning microscope (9–12 μm Z-stack with 0.5 μm step) magnified with 63x objective and first converted to projection images (with maximal projection) for analyses. The software SynPAnal 2 was used for quantifying the puncta density and intensity/area of PSD95^+^ and VGLUT1^+^ puncta. Neurite segments (20–30 μm in length) were quantified from each neuron and their average values were also measured using SynPAnal software.

For extracellular and intracellular CD63-GFP^+^ puncta analysis, the extracellular percentage ratio of CD63-GFP^+^ puncta were determined in relation to the tdT^+^ astroglia using Fiji ImageJ based on confocal images. The CD63-GFP channel image was first thresholded to create a binary black and red image. Then the Measure Analyzer tool was used to count all CD63-GFP^+^ puncta area. The tdT channel image was thresholded and the Particle Analyzer tool was used to generate the ROIs of all tdT^+^ signals. Then the ROIs of tdT^+^ signals were overlaid on the CD63-GFP^+^ images. CD63-GFP^+^ area was then measured inside of tdT based ROIs. CD63-GFP^+^ puncta inside tdT^+^ ROIs were considered as intracellular CD63-GFP^+^ signals. Extracellular CD63-GFP^+^ area was determined by subtracting CD63-GFP^+^ intracellular area from total CD63-GFP^+^ area and the extracellular percentage ratio was calculated by dividing the total CD63-GFP^+^ area by the extracellular CD63^+^ area.

For quantification of dendritic spine density, confocal images of eGFP^+^ pyramidal neurons of layer V motor cortex of Thy1-YFP^+^ and Thy1-YFP^+^ApoE^-/-^ mice were acquired at 0.5 μm intervals with a 63×oil immersion lens with Leica falcon confocal microscope. 3D reconstruction of eGFP^+^ neurons was built using the Imaris image analysis software (Bitplane). Both apical collateral and basal dendrites and spines were traced with the filament tracing function in Imaris and quantified. The dendritic spine density was calculated by dividing the number of spines by dendrite length (∼30 to 40 μm).

### *In vivo* anterograde labeling of CST axons and measurement of spinal cord CST axon length

CST axons were anterogradely labeled by a single injection of the CM-DiI dye (10 mg/mL in N, N-dimethylformamide) into the right-side motor cortex of P1 pups with the use of Hamilton micro syringe with 33 gage 30° needle. Pups were perfused at P3 with cold 1x PBS, brains with spinal cord were fixed in 4% PFA overnight, and 100μm sagittal cryosections were prepared along the anterior-posterior axis. They were mounted with Fluorogold anti-fade mounting medium then imaged under Keyence fluorescence microscope BZ-X700 with a Cy3 filter. Spinal cord CST axon length was measured based on the CM-DiI fluorescence signals from the superimposed images of individual mice (as shown in Fig. 5c) after the pyramidal decussation (PD) by using the segment line tool in ImageJ.

### Statistical analysis

All statistical analyses were performed and graphs were generated using GraphPad Prism 9. Group differences in each assay at each time point were analyzed by two-tailed t-test (2 group comparison), one-way ANOVA (3 or more group comparison, 1 independent variable), or two-way ANOVA (3 or more group comparison, 2 independent variables). Statistical test(s) used are specified in figure legends. Data are presented as mean ± SEM unless otherwise described. No custom code was used in the analysis. Statistical significance was tested at a 95% (p < 0.05) confidence level and p values are shown in each graph.

**Supplementary Fig. 1.**
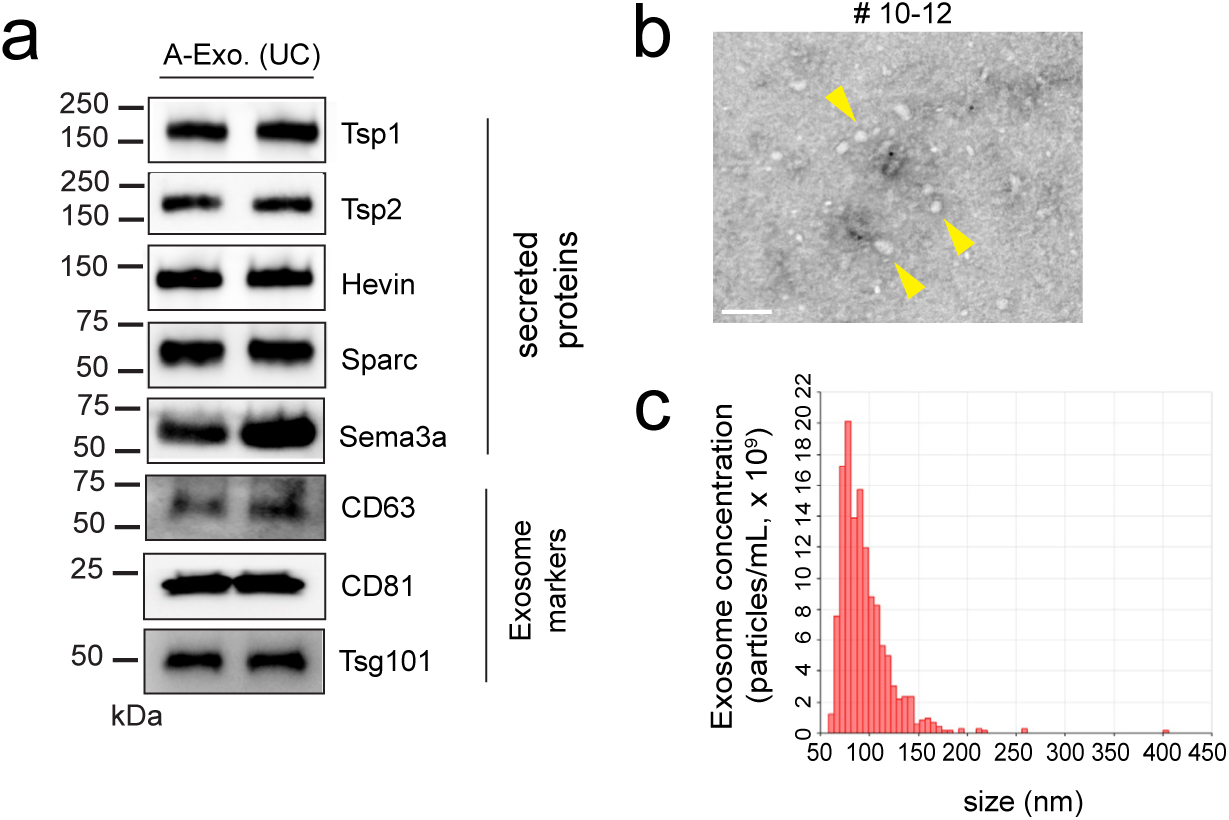
**a**, Representative immunoblots of astroglia secreted proteins and exosome markers in A-Exo. isolated from ACM (20 mL/sample) by ultracentrifυγation (UC). **b,** Representative immunoEM images of CD63 labeling in SEC eluted fractions #10-12; yellow arrows: CD63^-^ small vesicles; scale bar: 100 nm. **c**, Representative size distribution analysis of WT A-Exo measured by the qNano particle analyzer.

**Supplementary Fig. 2.**
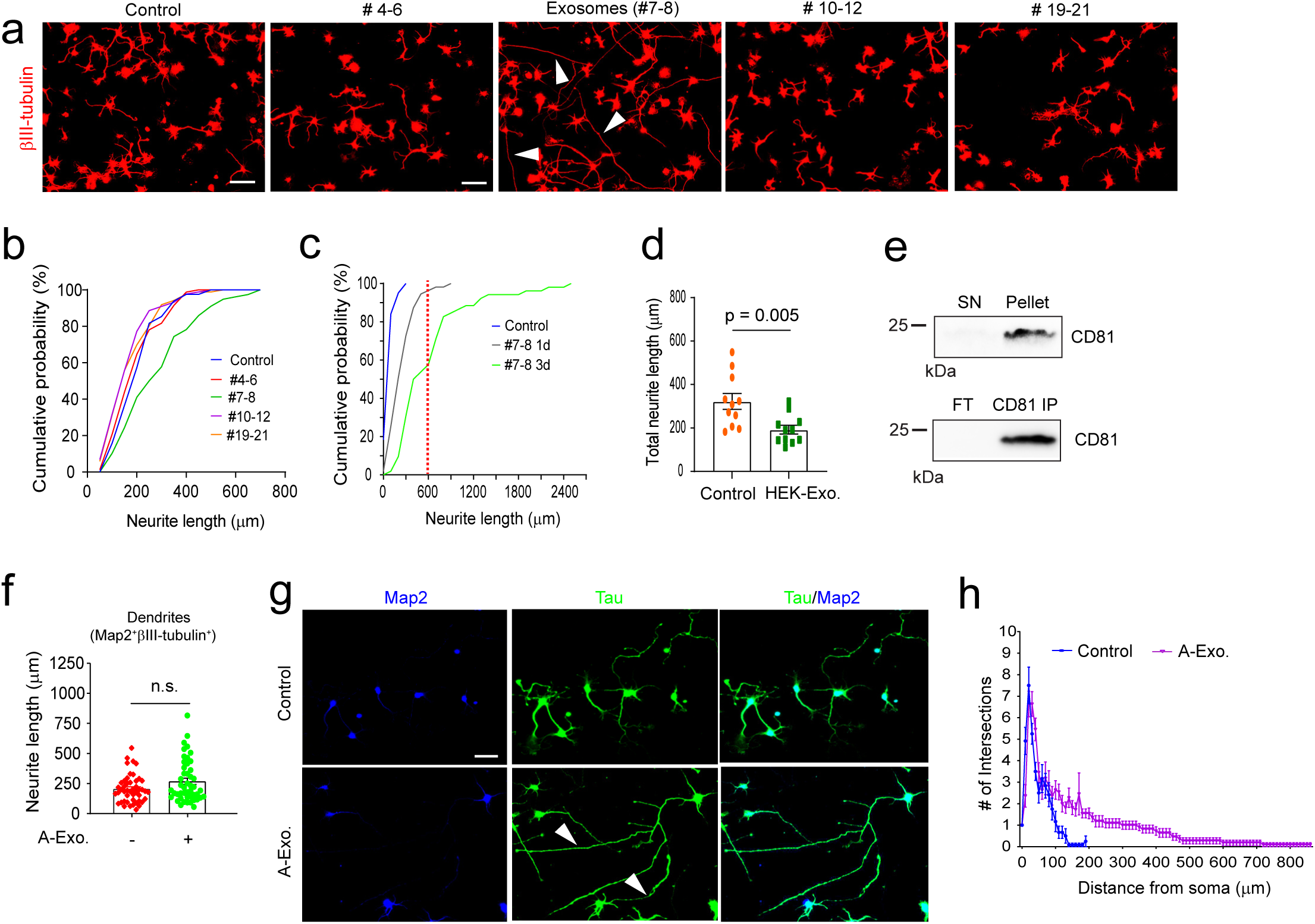
**a**, Representative images of βIII-tubulin^+^ cortical neurons following treatment with eluted fractions (pooled as indicated) #4-6, #7-8, #10-12, or #19-21 from the SEC. Scale bar: 100 μm; **b**, Quantification of total neurite length of cortical neurons following treatment with eluted fractions (pooled as indicated, 100 μl). #4-6 (no protein detected), #10-12, and #19-21 (1μg/μl) from the SEC of ACM (initial 100 mL). 1μg exosomes (#7-8) were used in treatment, n = 78-88 neurons (> 3 biological replicates)/group; **c**, Quantification of total neurite length of cortical neurons following treatment with fractions #7-8 (5μl, 0.2 μg/μl) for 1 or 3 d. n = 52-82 neurons (> 3 biological replicates)/group; **d**, Quantification of total neurite length of cortical neurons following treatment with HEK exosomes isolated by SEC. n = 11-13 neurons (2 biological replicates)/group; **e**, Representative immunoblot of CD81 in the supernatant (SN) or pellet of SEC fractions #7-8 (1 mL, from initial 10 mL ACM) following an additional 24 h ultracentrifugation (UC, 100,000 x g), or in the flowthrough (FT) or CD81 immunoprecipitation (IP) pellet of SEC fractions #7-8 after CD81 pull-down. **f**, Quantification of dendrite (Map2^+^βIII-tubulin^+^) length of cortical neurons following A-Exo. treatment. n = 51-55 neurons (> 3 biological replicates)/group; **g**, Representative images of Map2 and Tau staining on cortical neurons following A-Exo treatment. Scale bar: 50 μm; **h**, Sholl analysis of cortical neurons following A-Exo treatment. n = 10 neurons (2 biological replicates)/group; 1 μg exosome was used in **b-c, f,** and **h**. p values in **d** and **f** determined from two-tailed t test.

**Supplementary Fig. 3.**
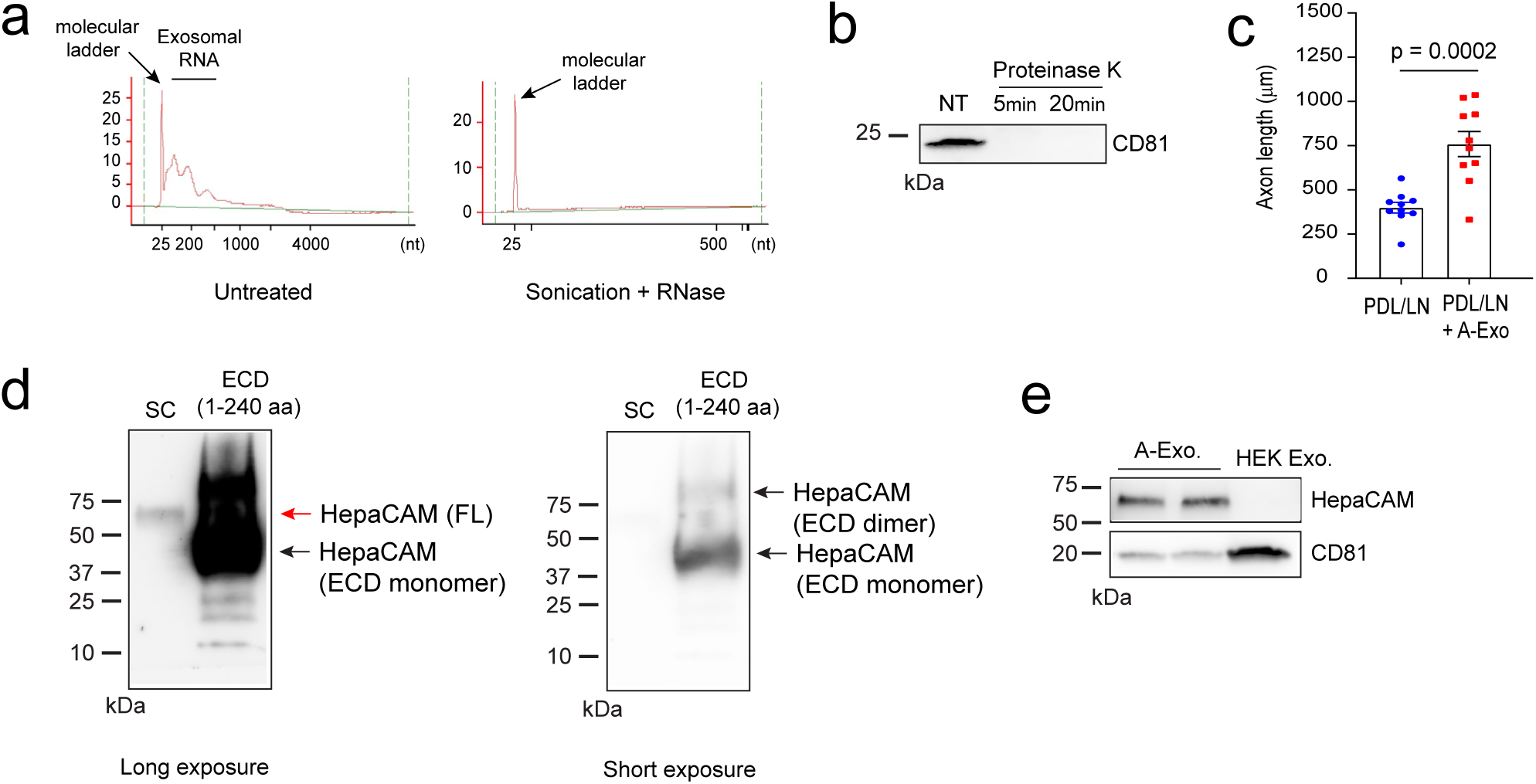
**a**, Representative bioanalyzer tracer of exosomal RNA with and without RNase treatment (5 minutes) following sonication. Sufficient small RNA was observed in untreated A-Exo. **b**, Representative immunoblot of CD81 following proteinase K treatment. 0.5 μg A-Exo. was treated with proteinase K for either 5 or 20 minutes. NT: not treated A-Exo; CD81 immunoreactivity disappeared from the immunoblot as a result of the proteinase K digestion; **c**, Quantification of axon length of cortical neurons plated on either PDL/laminin (LN) coated or PDL/LN/A-Exo. coated coverslips. n = 10 neurons (2 biological replicates)/group; 1μg A-Exo. was used in each treatment. p value in **c** determined from two-tailed t test; **d**, Representative HepaCAM immunoblot with spinal cord (sc) lysate (20 μg) and recombinant human HepaCAM extracellular domain (ECD) protein (1-240 aa, 1 μg). HepaCAM antibody (Proteintech) is able to detect mouse HepaCAM full-length (sc lane) and human ECD (monomer and dimer). **e**, Representative HepaCAM immunoblot in A-Exo. and HEK exosomes.

**Supplementary Fig. 4.**
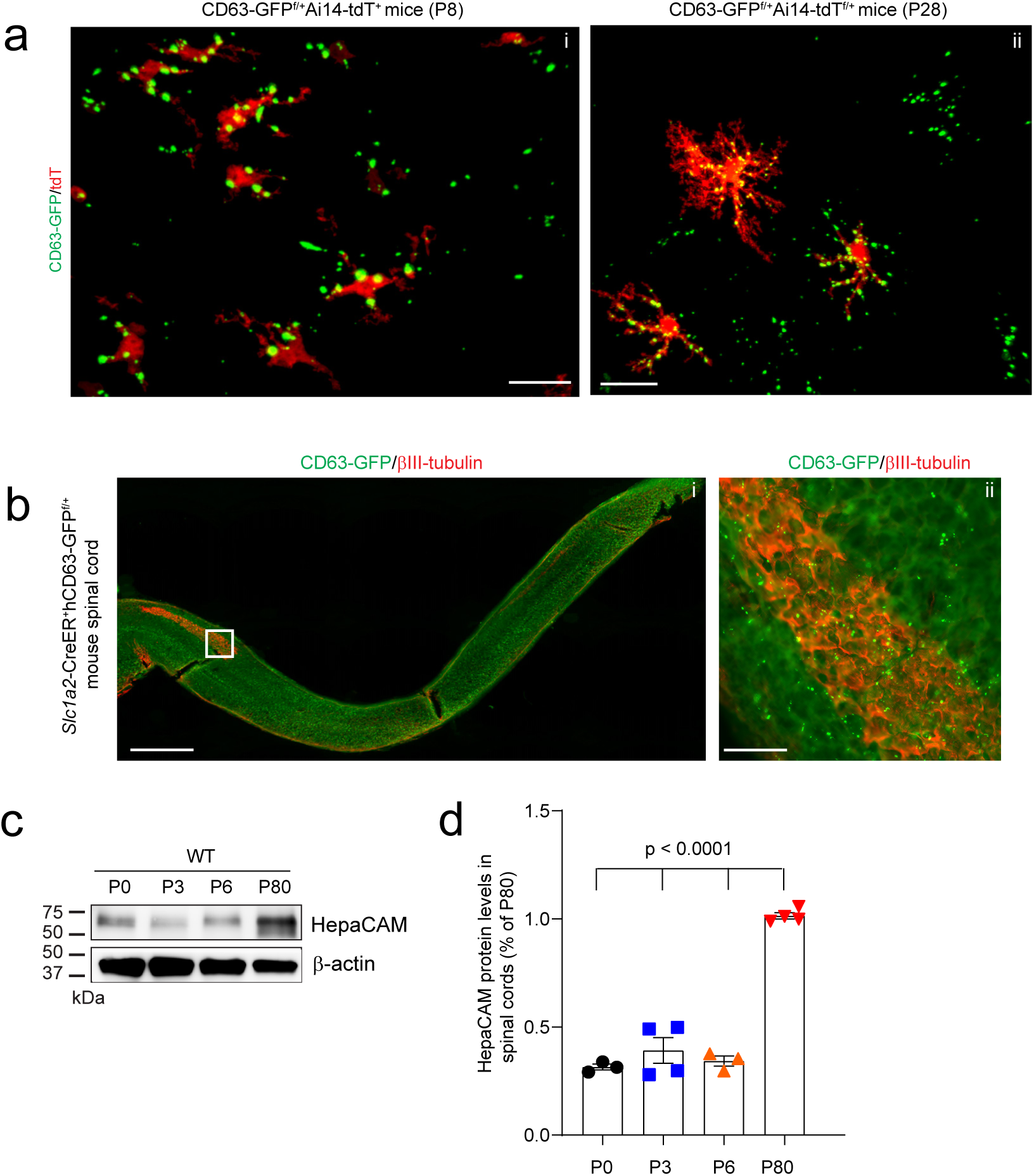
**a**, Representative images of tdT^+^ astroglia and astroglia-derived CD63-GFP^+^ puncta from the motor cortex of AAV5-mCherry-*Gfap*-Cre-injected CD63-GFP^f/+^Ai14-tdT^f/+^ mice at P8 (i) and P28 (ii). Scale bar: 20 μm; **b**, Representative longitudinal image of βIII-tubulin staining and astroglia-δεριϖεδ CD63-GFP^+^ puncta along the spinal cord from 4-OHT-injected *Slc1a3*-CreER^+^ mice at P8. Subpanel i: the longitudinal image of the spinal cord; Subpanel ii: a magnified view of the box in the subpanel i; Scale bar: 1mm (subpanel i); 100 μm (subpanel ii); Representative HepaCAM immunoblot (**c**) and quantification (**d**) of HepaCAM expression in spinal cords during postnatal development. n = 3-4 mice/time point; p values determined by one-way ANOVA followed by post-hoc Tukey’s test.

**Supplementary Fig. 5.**
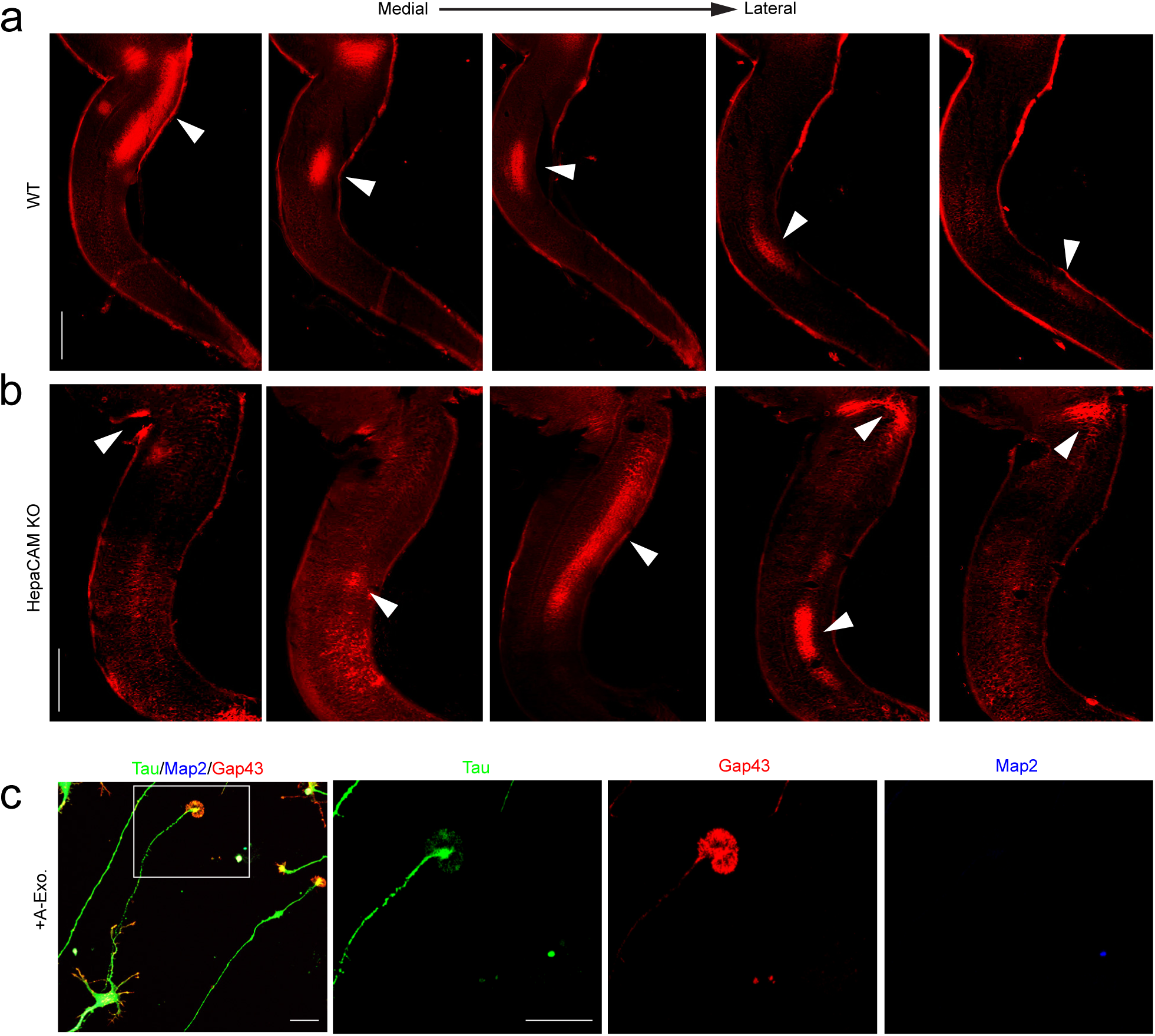
Representative original set of longitudinal images from CM-DiI-injected WT (**a**) and HepaCAM KO (**b**) mouse spinal cords that were superimposed into the continuous CST axon growth image shown in Fig. 5C. Images of longitudinal spinal cord sections were taken from lateral to medial orientation at P3. White arrows: CM-DiI labeling; Scale bar: 1mm; **c**, Representative image of Tau, Map2, and Gap43 immunostaining of A-Exo-treated cultured cortical neurons to illustrate axonal growth cones and axons; Scale bar: 20 µm.

**Supplementary Fig. 6.**
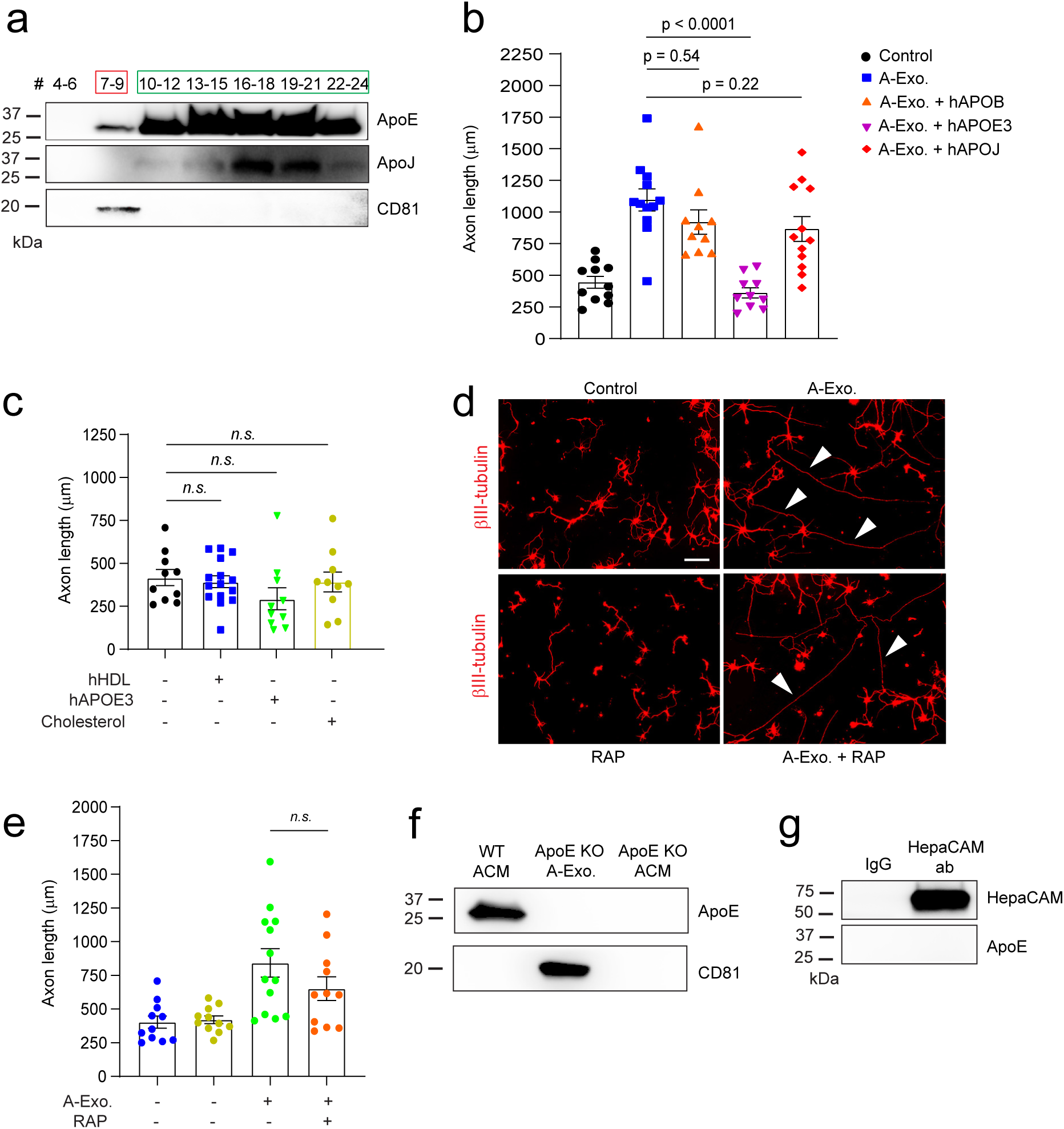
**a**, Representative immunoblot of ApoE and ApoJ in all eluted fractions (500 μl/fraction, pooled as indicated) of ACM (100 mL) from SEC with oversaturated exposure. 15 μl unconcentrated elution was run on immunoblot. **b**, Quantification of βIII-tubulin^+^ neuronal axon length following co-treatment of A-Exo. with hAPOEB, hAPOEJ, or hAPOE3, respectively. 1μg A-Exo. was used in treatment. hAPOEB, hAPOEJ, or hAPOE3 each was at 10 μg/mL dose. n = 11-12 neurons (2 biological replicates)/group; **c**, Quantification of βIII-tubulin^+^ neuronal axon length following treatment of hHDL (10 μg/mL), hApoE3 (20 μg/mL), and cholesterol (1 μg/mL), respectively. n = 10-15 neurons (2 biological replicates)/group; Representative images (**d**) and quantification (**e**) of βIII-tubulin^+^ neuronal axon (white arrows) length following co-treatment of A-Exo. and ApoE competitive receptor associated protein (RAP, 50 μg/mL). Scale bar: 100 μm; **f**, Representative ApoE immunoblot from WT or ApoE ACM (50μg proteins), and ApoE A-Exo (2 μg proteins). **g**, Detection of HepaCAM but not Apoe following HepaCAM immunoprecipitation from astrocyte lysates (50 μg proteins). 1μg A-Exo. was used in **b, d**, **e,** and **f**. p values in **b, c,** and **e** determined from one-way ANOVA followed by a Tukey post-hoc test.

**Supplementary Fig. 7.**
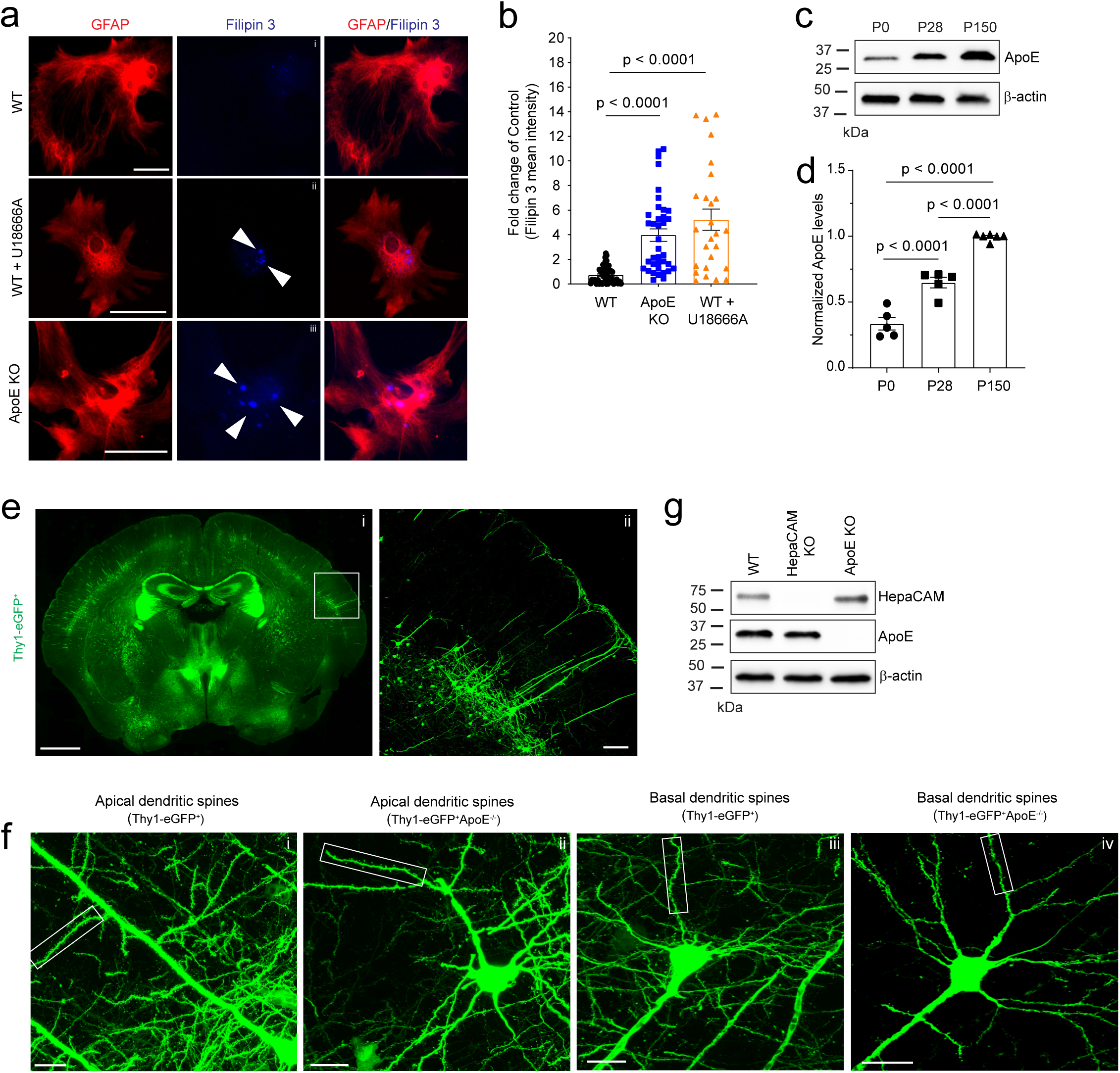
Representative images (**a**) of cultured astrocytes and cholesterol labeling and quantification (**b**) of cholesterol in astrocytes based on Filipin 3 fluorescent intensity. Scale bar: 50 μm; White arrows: Filipin 3^+^ cholesterol labeling; n = 26-35 astrocytes (3 biological replicates)/group; Representative images (**c**) of ApoE immunoblot and quantification (**d**) of ApoE expression in the cortex during postnatal development; n = 5-6 mice/group; **e**, eGFP labeling of neurons and neurites in Thy1-eGFP^+^ mice. Subpanel i: Representative image of coronal section of the Thy1-eGFP^+^ mouse brain (scale bar: 1mm); ii: a magnified view of the motor cortex (white box) in the subpanel i (scale bar: 100 μm); **f**, Representative images of eGFP^+^ neurons and their dendritic spines. Subpanel i: apical dendritic spines from Thy1-eGFP^+^ mice; ii: apical dendritic spines from Thy1-eGFP^+^ApoE^-/-^ mice; iii: basal dendritic spines from Thy1-eGFP^+^ mice; iv: basal dendritic spines from Thy1-eGFP^+^ApoE^-/-^ mice; Scale bars: 20 μm; a magnified view of the highlighted box is shown in Fig. 7d-e; **g**, Representative HepaCAM and ApoE immunoblots from cortex of ApoE KO and HepaCAM KO mice at P30.

**Supplementary Table 1.** Transmembrane proteins identified from A-Exo. by LC/MS/MS. Each identified protein has at least 3 peptide hits with 95% confidence threshold; The mean iBAQ value is greater than 1 x 10^5^.

**Supplementary Movies** Live imaging of control and A-Exo (1μg). -induced axon growth in primary cortical neuronal cultures.

